# Neuronal Spike Shapes (NSS): A Straightforward Approach to Investigate Heterogeneity in Neuronal Excitability States

**DOI:** 10.1101/2023.06.28.546812

**Authors:** Lorenzo Martini, Gianluca Amprimo, Stefano Di Carlo, Gabriella Olmo, Claudia Ferraris, Alessandro Savino, Roberta Bardini

**Author notes:** Corresponding author (L. Martini); (R. Bardini). Principal corresponding author (G. Amprimo). These authors share co-first authorship and contributed equally to this work (Lorenzo Martini, Gianluca Amprimo). www.smilies.polito.it (L. Martini); https://www.polito.it/personale?p=gianluca.amprimo (G. Amprimo); www.smilies.polito.it (S. Di Carlo); https://www.sysbio.polito.it/analytics-technologies-health/ (G. Olmo); https://www.ieiit.cnr.it/people/Ferraris-Claudia (C. Ferraris); www.smilies.polito.it (A. Savino); www.smilies.polito.it (R. Bardini).

## Abstract

The mammalian brain exhibits a remarkable diversity of neurons, contributing to its intricate architecture and functional complexity. The analysis of multimodal single-cell datasets enables the investigation of cell types and states heterogeneity. In this study, we introduce the Neuronal Spike Shapes (NSS), a straightforward approach for the exploration of excitability states of neurons based on their Action Potential (AP) waveforms. The NSS method describes the AP waveform based on a triangular representation complemented by a set of derived electrophysiological (EP) features. To support this hypothesis, we validate the proposed approach on two datasets of murine cortical neurons, focusing it on GABAergic neurons. The validation process involves a combination of NSS-based clustering analysis, features exploration, Differential Expression (DE), and Gene Ontology (GO) enrichment analysis. Results show that the NSS-based analysis captures neuronal excitability states that possess biological relevance independently of cell subtype. In particular, Neuronal Spike Shapes (NSS) captures, among others, a well-characterized fast-spiking excitability state, supported by both electrophysiological and transcriptomic validation. Gene Ontology Enrichment Analysis reveals voltage-gated potassium (*K*^+^) channels as specific markers of the identified NSS partitions. This finding strongly corroborates the biological relevance of NSS partitions as excitability states, as the expression of voltage-gated *K*^+^ channels regulates the hyperpolarization phase of the AP, being directly implicated in the regulation of neuronal excitability.

## 1. Introduction

The mammalian brain displays many cell types, contributing to its complex structure and functions [30, 1, 72, 16]. Investigating cellular heterogeneity means understanding how cells specialize for specific functions [43, 54] and how they interact to form intricate neuronal networks during development or in adult organisms [20, 14, 74, 36, 37].

Multimodal single-cell analysis provides multiple layers of data to explore neuronal states and their transcriptional profiles across cell types [47, 46, 28, 27, 78]. Indeed, understanding cellular heterogeneity requires profiling thousands of cells in multiple dimensions to accurately identify and characterize neuronal types and subtypes [80, 59, 8].

A cell type is a group of cells that share a stable phenotype based on transcriptomic, morphological, and functional markers. However, these categories have significant uncertainties [84]. Neurons exhibit a remarkable diversity in terms of their types. Neuronal types encompass various categories, each classified according to distinct sets of criteria, often overlapping. This classification can hinge on morphological features, such as the neuron’s shape, whether it resembles a pyramid, star, or exhibits a bipolar structure. Alternatively, neuronal categorization can be based on their functional roles within neural circuits or the specific neurotransmitters they release. In the context of this study, our primary focus is on GABAergic neurons, which release the inhibitory neurotransmitter GABA (gamma-aminobutyric acid) and play a crucial role in modulating the activity of excitatory neurons. It is worth noting that GABAergic neurons comprise multiple cell subtypes, each characterized by distinct morphological and transcriptomic profiles.

Furthermore, the varying nature of cell dynamics introduces additional layers of heterogeneity, introducing the concept of cell states. A cell state refers to a *“transient or dynamically responsive property of a cell to a context”* [84], distinguished by complex molecular and functional properties [55]. These states encompass varying activation levels and involve neuroplasticity mechanisms, which are connected to the dynamic adaptation of neurons to activation patterns within a neural circuit [12]. In the mammalian brain, plasticity mechanisms play a crucial role in the variability of neuronal states, acting throughout the lifespan of the organism to balance stability and adaptation of neuronal networks, supporting brain functions and learning [77, 34].

Investigating these states is paramount, as it is crucial to understand the intricate biological mechanisms underlying brain functions [50]. However, the identification of cell states is challenging due to the lack of well-defined, quantitative, and precise state definitions in the literature. This results in a notable gap concerning dedicated methods for effectively examining these states.

This paper introduces Neuronal Spike Shapes (NSS), which represents an innovative model-based approach crafted to explore electrophysiological data to study cell states within neurons. To bridge the existing gap in the investigation of excitability states, NSS introduces a simple model that requires a limited amount of electrophysiological (EP) data, summarizing the Action Potential (AP) waveform into a triangular representation through a set of descriptive EP features. What sets NSS apart from existing research on cell heterogeneity studies is that NSS does not operate as a classification method aimed at assigning labels to cell types, thereby constructing a comprehensive taxonomy encompassing various neuronal categories. Instead, NSS adopts a structured approach to perform unsupervised exploratory analysis, aiming to identify electrophysiological cell states within groups of cells. In addition, using only AP-related features among the many other EP features, the provided approach requires a minimal set of data compared to other approaches, mitigating data availability issues that are critical for patch-clamp and in particular patch-seq datasets. It is worth noting that the current literature still lacks dedicated methods for effectively probing cell states at the single-cell level, underscoring the significance of our work.

This study applies the NSS analysis to two high-throughput datasets of murine cortical interneurons. The first dataset provides solely electrophysiological data [25], while the second dataset offers a comprehensive characterization of cells encompassing electrophysiological, transcriptomic, and morphological aspects [24]. Results validate NSS’s capacity to autonomously identify well-established cell states in neurons, outperforming existing approaches detailed in the literature. Given the limited body of related work, our validation primarily concentrates on identifying the high excitability fast-spiking state in neurons [23]. Results demonstrate NSS’s capability for identifying cells within high excitability state by comprehensively evaluating different AP features. Furthermore, our approach successfully identifies new clusters of cells that share AP features, thereby representing unknown excitability states, thus providing a guide for future investigations. This outcome underscores the exploratory potential of the proposed pipeline, which can unveil concealed relationships among states within a dataset. These relationships can subsequently be further examined in conjunction with cell types to advance our comprehension of neuronal behavior.

After providing an overview of the scientific background in Section 2 and of the existing literature in Section 3, this manuscript presents the proposed methodology in Section 4, followed by a detailed presentation and discussion of the conducted analysis and its results in Section 5. Section 6 draws conclusions and suggests future directions to advance the proposed research.

## 2. Background

Current approaches to studying EP characteristics related to cell states involve the computational analysis of single-cell multimodal data and the development of mathematical models to describe the AP waveform. This section introduces essential background concepts for comprehending the proposed methodology and its relevance to the study of neuronal states.

### 2.1. Computational electrophysiological analysis of neuronal states

Computational studies focusing on cell heterogeneity aim to uncover distinct cell heterogeneity within a single-cell dataset [51, 81, 32, 61, 25, 24, 39]. Typically, this involves conducting unsupervised clustering analysis to group cells and assigning identities to these clusters based on established knowledge [8]. For example, in the context of single cell RNA sequencing (scRNA-seq) data, cells are labeled by identifying known markers within their expression profiles [33, 79]. However, this marker-based approach prioritizes recognizing stable cell identities over detecting transient functional states [53, 32]. In this context, the use of multimodal datasets provides a valuable opportunity to enhance the study of cell heterogeneity and state dynamics by integrating consistent information from multiple modalities obtained from the same cells [27].

Considering state dynamics in the multimodal analysis of neuronal heterogeneity is pivotal. Indeed, information processing in neuronal cells relies both on the characteristics of the stimulus, such as its intensity and on transient neuronal cell states, such as excitability states [71]. Constantly shifting excitability states make a neuron’s reaction to the same stimulus vary from one moment to another and affect functional activation [45]. Thus, the regulation of membrane excitability is a fundamental physiological phenomenon that drives the operation of all biological tissues in which voltage-dependent changes in ionic conductances lead to APs [45]. In the brain, activity-dependent modulation of intrinsic excitability plays a significant role in the plasticity of neuronal circuits [73]. The AP waveform comprises extrinsic components resulting from interactions between neurons and other cells, as well as the extracellular environment. It also involves dynamic regulation of membrane composition driven by neuroplasticity mechanisms that operate on different timescales[18, 50]. These factors transiently affect the neuronal excitability state at the time of stimulation.

Patch-seq, a multimodal single-cell technology, holds particular significance in studying neuronal cell states as it simultaneously analyzes electrophysiological activity and transcriptomic profile [49, 42, 10, 9]. In fact, by directly accessing data on the functional cellular responses to defined stimuli, patch-seq supports the analysis of cellular activation patterns as markers of transient functional states. The electrophysiological component of Patch-seq relies on the patchclamp technique for studying ion channels [56, 57, 29]. This enables the characterization of cellular electrophysiological properties and provides a platform for basic and pharmacological research [19, 41, 58]. During a Patch-seq analysis, cells exhibit a range of electrophysiological responses. Among them, the NSS approach specifically focuses on the action potential waveform due to its ability to provide crucial information on the neuronal excitability state [45].

### 2.2. The action potential waveform

In neurons, the Action Potential waveform provides information on the excitability state. APs mediate cell-cell communication, allowing signals to propagate along the axon and reach synaptic buttons at the axon terminals [3]. An AP is characterized by a rapid sequence of voltage changes that occur across the neuronal membrane [4]. Over time, the membrane potential is determined by the relative ratio of extracellular and intracellular ion concentrations and the membrane permeability to each ion. Although the AP waveform remains consistent between species and neuronal cells, differences in the composition of ion channel populations in the neuronal membrane lead to noticeable variations in AP shape, relating to the cell’s excitability state. These dynamics follow well-conserved phases, as described in this section.

The AP can be divided into four primary phases, as illustrated in Figure 1:

**Figure 1:**
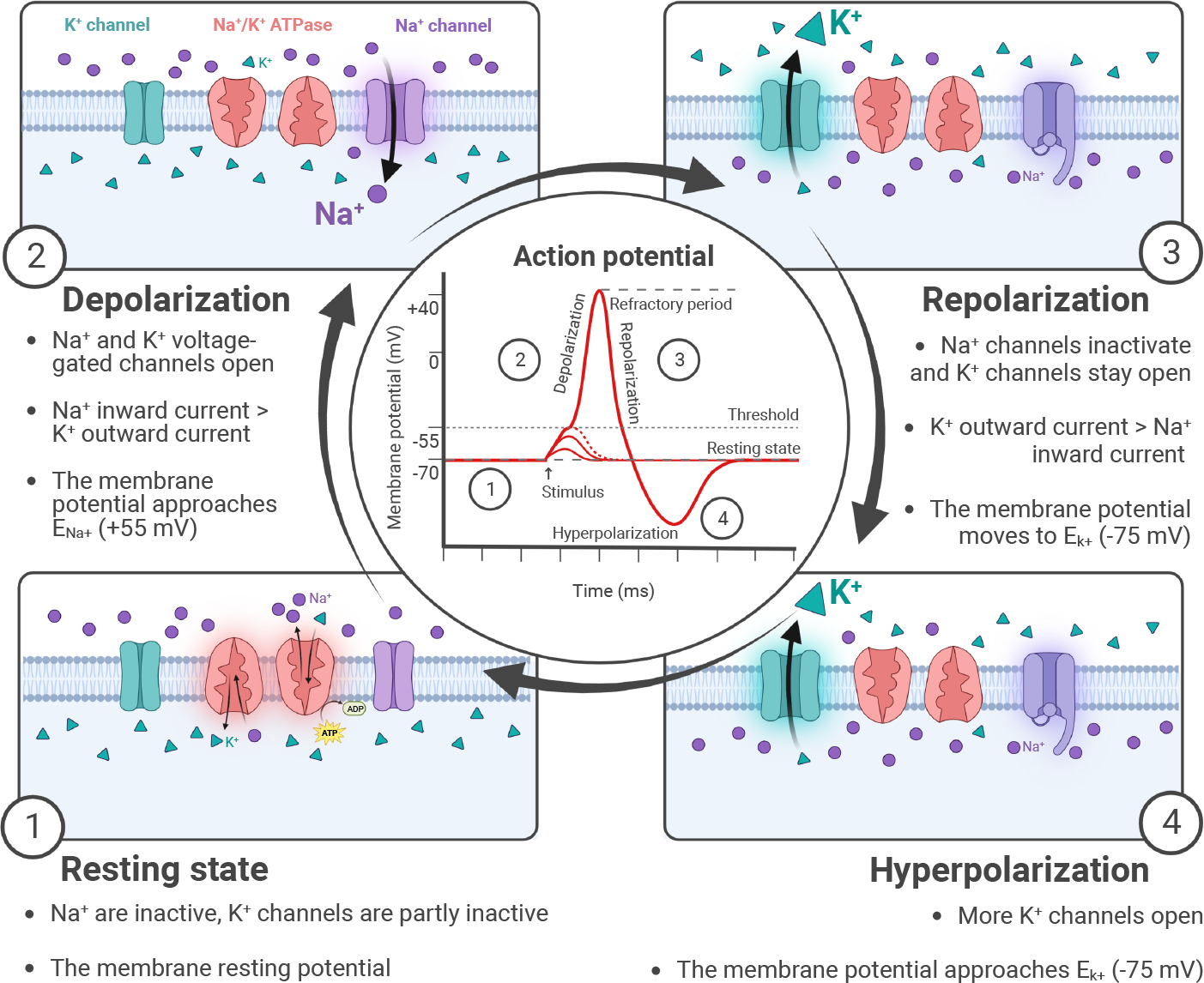
Membrane dynamics underlying AP. (1) Resting state. The membrane resting potential is close to the K_+_ equilibrium potential 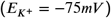, and at this potential both Na_+_ and K_+_ voltage-gated channels are closed. (2) Depolarization. When the AP is triggered, membrane depolarization causes the voltage-gated Na_+_ and K_+_ channels to open. It starts positive feedback where depolarization causes the opening of more Na_+_ channels, which in turn further depolarizes the cell membrane until reaching a positive value that approaches the Na_+_ equilibrium potential 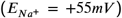 (3) Repolarization. As membrane potential gains positive values, Na_+_ channels begin to inactivate, and K_+_ channels stay open, stopping the Na_+_ ions flux into the cell, and making the flow of K+ ions outside the cell prevail (4) Hyperpolarization. Na_+_ channels are still inactivated, and more K_+_ channels open due to repolarization and the Ca_2+_ influx caused by the AP. Na_+_ channels inactivation causes a refractory period where subsequent stimuli can not trigger APs. Then, the membrane slowly returns to its resting potential.

1. **Resting state:** In this phase, the membrane potential is close to the equilibrium potential of potassium ions 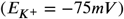, and both sodium (*Na*^+^) and potassium (*K*^+^) voltage-gated channels are closed (Figure 1.1).
2. **Depolarization:** When the AP is triggered, depolarization of the membrane occurs, resulting in the opening of voltage-gated *Na*^+^ and *K*^+^ channels. This triggers a positive feedback loop, where depolarization opens more *Na*^+^ channels, further increasing membrane depolarization. The membrane potential approaches a positive value that is closer to the equilibrium potential of sodium ions 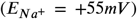 (Figure 1.2).
3. **Repolarization:** As the membrane potential becomes positive, *Na*^+^ channels begin to inactivate, while *K*^+^ channels remain open. This halts the influx of *Na*^+^ ions into the cell and allows the efflux of *K*^+^ ions to prevail, leading to repolarization (Figure 1.3).
4. **Hyperpolarization:** During this phase, *Na*^+^ channels remain inactivated and the repolarization process causes additional *K*^+^ channels to open. Furthermore, the AP triggers the influx of calcium ions (*Ca*^2+^). As a result, the membrane potential briefly becomes more negative than the resting state (Figure 1.4).

**Figure 2:**
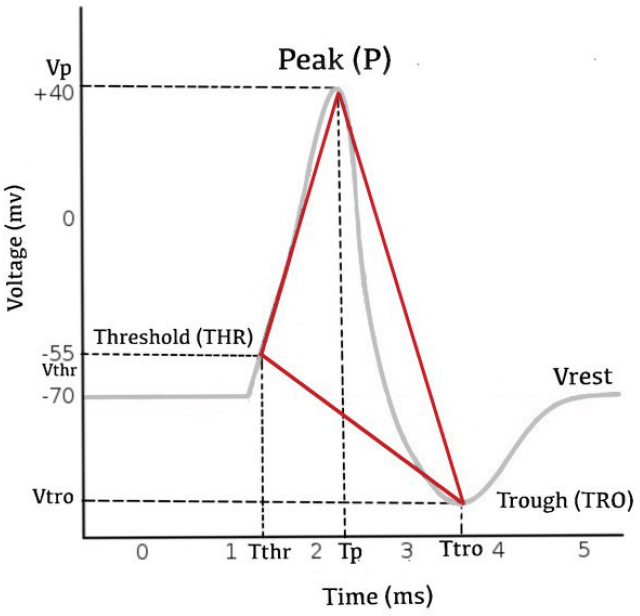
Example of neuronal AP. Threshold (THR), Peak (P), and Trough (TRO) points are the vertices of the NSS triangular representation (in red) over-imposed to the AP waveform.

**Figure 3:**
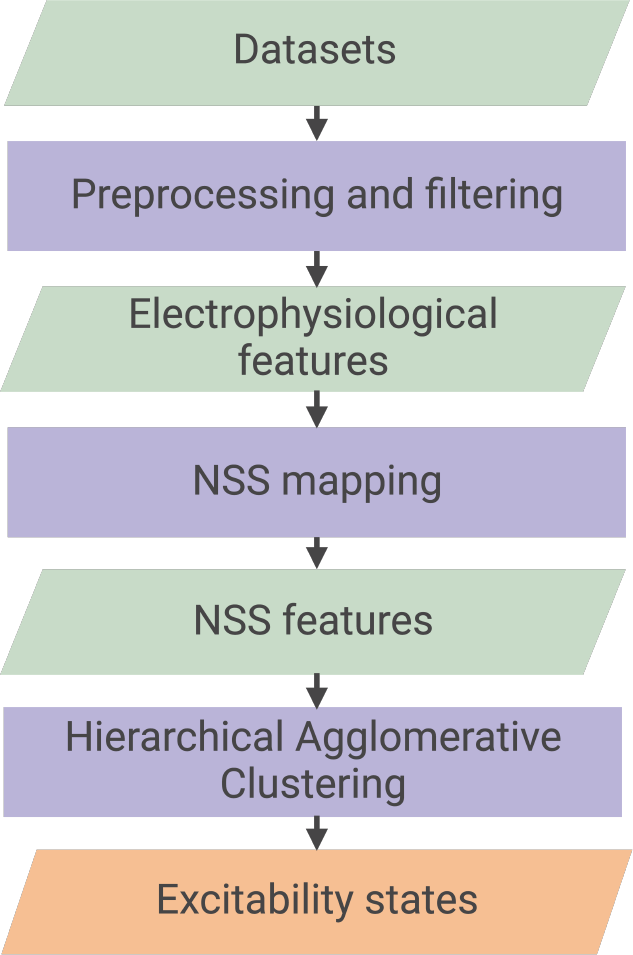
Pipeline of EP exploration using NSS on*PatchClampDataset* and *PatchSeqDataset*.

**Figure 4:**
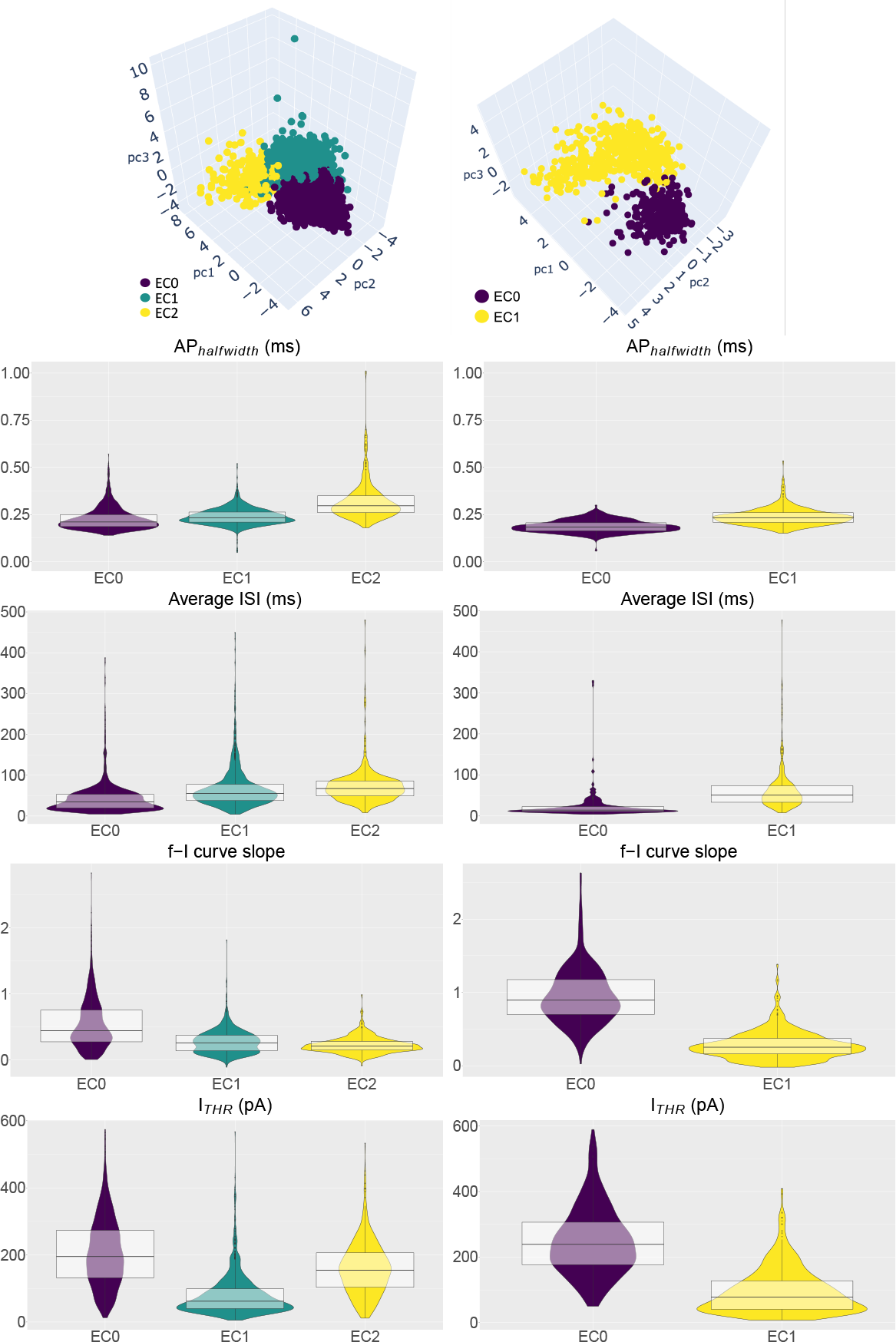
NSS clustering results and fast-spiking EP feature analysis for *PatchSeqDataset* (right column) and *PatchClampDataset (left column)*. In the first row, tri-dimensional PCA plots along the three most relevant PCs visualize clustering results for *PatchSeqDataset* (*K* = 3) and *PatchClampDataset* (*K* = 2). The following rows present violin and box plots of the distribution of cells within each NSS cluster over fast-spiking feature values The color coding refers to clustering labels: for *PatchSeqDataset* (*K* = 3) *EC0* (violet), *EC1* (green), and *EC2* (yellow), for *PatchClampDataset* (*K* = 2) *EC0* (violet) and *EC1* (yellow).

Following the peak of the AP, the voltage-gated *Na*^+^ channels enter an inactivated state, resulting in a refractory period where subsequent stimuli can not trigger APs. Over time, the *Na*^+^ channels gradually reactivate, while the *Na*^+^*K*^+^ ion pump actively restores the resting membrane potential.

### 2.3. Models of the action potential waveform

The initial model of the AP, as proposed by Hodgkin and Huxley, described its initiation and propagation in neuronal cells using a set of nonlinear differential equations, representing a continuous-time dynamical system [31]. However, this model considered only *Na*^+^ and *K*^+^ conductances, each with a single type of voltage-dependent channel. This simplistic approach overlooked the complexity of voltage-dependent conductances in neuronal membranes, limiting the exploration of the underlying biomolecular substrates of electrophysiological behavior. In reality, the neuronal membrane exhibits a wide diversity of conductances. Voltagegated currents, including *Na*^+^, *Ca*^2+^, and *K*^+^ currents, consist of at least two distinct components [4]. Additionally, other currents are activated by membrane hyperpolarization [6]. The complexity and diversity of AP conductances can be attributed to a wide range of types and isoforms of ion channels [35, 82]. The relative prevalence of different channel populations at the membrane contributes to the electrophysiological features of neuronal action potentials. The interplay and regulation of channel activation among these populations present a significant challenge in understanding and modeling this phenomenon [4]. Alterations in the underlying biomolecular ion channel composition impact AP shape, firing rate, and the emergence of distinct higher-level patterns in the electrophysiological phenotype and state of different neurons [4].

The intrinsic components of the AP waveform generated by a particular neuron are regulated by its electrophysiological and morphological features and its membership in specific groups of cells (as demonstrated by the taxonomy described in [24]). In fact, the AP waveform exhibits substantial and stable differences among various neurons in the mammalian brain [4]. Consequently, the analysis of AP waveforms alone facilitates the study of cellular heterogeneity among interneuronal cell families [13]. Mathematical models of AP shape have supported the construction of a coherent neuronal taxonomy that aligns with the transcriptomic profiles of cells [63].

Given the intricate structural and functional interconnections among neurons within brain neuronal networks, each AP waveform property is intrinsic, dependent on the cell itself, and extrinsic, influenced by stimulation and cell connections during stimulation. Consequently, the AP waveform provides information on cell states and the corresponding functional activations of neurons within their respective neuronal networks. The proposed approach uses the rich information contained in the AP waveforms to analyze neuronal excitability states.

## 3. Literature review

While the literature provides a rich set of approaches for classifying cell types using electrophysiological features [25, 24] [63], it lacks approaches specifically addressing cell states, which is the primary focus of NSS. Indeed, the literature lacks a clear and quantitative definition of intrinsic excitability states, making it challenging to understand their role in neuronal function and network dynamics. Furthermore, traditional methods to assess intrinsic excitability fail when exposed to realistic synaptic input [73].

Current approaches in studying intrinsic excitability states involve analyzing various EP features. Some methods focus on overall firing rates to assess excitability, assuming that a neuron’s average firing rate encodes information about its intrinsic excitability and input received [75]. For instance, research by [62] explores the effect of dopamine on intrinsic excitability states by measuring cortical interneuron firing rates (which range from about 8 Hz in control to reaching 70 Hz under stimulation). Other studies, such as [17], also employ spiking rates to gauge excitability. Some investigations concentrate on AP shape for excitability analysis. For example, [68] focuses on the AP repolarization phase duration, [13] and [40] base their analysis on AP total duration, while [22] defines fast-spiking neocortical interneurons’ excitability state using various AP characteristics.

These approaches collectively underscore the importance of EP features and AP shape in understanding intrinsic excitability. However, they reveal a lack of consensus over the EP features and their distinctive ranges for known excitability states. For example, several works measure intrinsic excitability based on the AP half-width, but describe the same excitability state, i.e., physiological fast-spiking behavior in neocortical interneurons, with different measures of AP half-width: 0.25 ± 0.02 ms in [40], 0.34 ± 0.07 ms in [22], and 0.64 ± 0.04 ms in [11]. These limitations highlight the need for a more structured definition of excitability.

From the reviewed literature, excitability definitions are either *activation-based, threshold-based*, or both. Activation-based excitability is based on the *intensity of the firing activity in relation to the stimulus*. Considering the latter, highly excitable cells have high firing rates [62] compared to other cells at the same level of stimulation. Threshold-based excitability is based on the *ease of evoking a AP response*. In this sense, highly excitable cells have low AP thresholds [40], implying a low-intensity current stimulus is sufficient to evoke a firing response. However, activation- and threshold-based definitions sometimes lack coherence. For example, fast-spiking neocortical interneurons are described as cells in a high excitability state, and present high firing rates (coherently with activation-based definitions) but high AP thresholds (contrary to what threshold-based definition would imply) [17, 23]. This example is especially relevant for the provided work, which aims to analyze two datasets of cortical interneurons.

To address the current lack of a unique and comprehensive definition of excitability, a more systematic and quantitative approach is required, involving intentional selection of EP features and dynamic range analysis rather than fixed, experiment-dependent values [45]. In fact, the NSS methodology provides an automatic, unsupervised quantitative pipeline to analyze neuronal excitability states, based primarily on the AP waveform. It supports the joint analysis of relevant features, enabling systematic exploration of excitability states in patch-seq and patch-clamp datasets. NSS offers the advantage of unsupervised cell grouping, automatically identifying states and their corresponding feature ranges, generalizing the method across datasets.

Additionally, NSS relies on a minimal set of EP features describing the first AP obtained during patch-clamp ramp stimulation. This minimizes the number of required experiments, making the method efficient and practical.

## 4. Methods

The NSS method utilizes a simplified triangular representation of the AP waveform (see Section 4.1), allowing a concise description of individual neuronal spikes. This triangular shape serves as a basis for computing mathematical electrophysiological features, which summarize the shape and duration of the AP generated by a neuronal cell.

Due to the relations between the AP shape and the firing pattern of neurons, NSS describes excitability states, corresponding to the ability of cells to respond with different intensities and firing patterns to stimuli, and not as the easiness to evoke an AP response.

### 4.1. NSS: a simple triangular model

The proposed mathematical model summarizes the attributes of a neuron’s AP waveform into a straightforward triangular representation within the voltage-time plane. The AP waveform exhibits a well-conserved spike shape in the time-voltage domain, as shown in Figure 2. Within the NSS framework, three points are considered particularly informative regarding the AP:

- **Threshold (THR)**: This point corresponds to the minimum depolarization of the cell membrane relative to the resting potential (see Figure 1.1) required to initiate an AP. The threshold dynamically adapts based on previous membrane activations [60] and varies depending on the cell type and state. For example, the voltage threshold for AP generation in cortical interneurons of the Fast-spiking (FS) and Non-Fastspiking (NFS) types was found to be approximately−31.6 ± 5.0 mV and −39.6 ± 6.4 mV, respectively [23].
- **Peak (P)**: This point corresponds to the highest depolarized voltage attained during AP generation (i.e., the peak of the spike). At the peak, the membrane exhibits maximum permeability to sodium ions (*Na*^+^), causing the membrane potential to approach the *Na*^+^ equilibrium potential 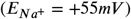. Peak potential typically ranges from 0 to +40 mV, as reported in [5] (see Figure 1.1).
- **Trough (TRO)**: This value, as defined in [25] and [24], represents the hyperpolarization phase (see Figure 1.4). It can be further divided into two sub-points: the *Fast Trough (FTRO)* corresponding to the most negative membrane potential within 5 ms after the peak time, and the *Slow Trough (STRO)*, corresponding to the most negative membrane potential between 5 ms after the peak time and the subsequent action potential threshold time. During the hyperpolarization phase, the membrane potential approaches the equilibrium potential of potassium ions (*K*^+^) 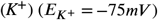, with values typically ranging from −70 to −75*mV*, as reported in [5].

This triangular representation of the AP serves as the fundamental framework of the NSS approach. The coordinates of its three vertices are determined by the voltage values (*V*_*T HR*_, *V*_*P*_, *V*_*F T RO*_) and time values (*T*_*T HR*_, *T*_*P*_, *T*_*F T RO*_) specific to each analyzed AP potential. Furthermore, the sides of the triangle summarize various functionally relevant geometric properties of the AP waveform. Specifically, the THR − P side approximates the depolarization phase (see Figure 1.2), the P − FTRO side represents the repolarization phase (see Figure 1.3) and hyperpolarization phase (see Figure 1.4), while the THR − FTRO side reflects the overall duration of the AP and the depth of the hyperpolarization.

### 4.2. NSS Features

Using the abstract triangular simplification at the core of the NSS approach, it becomes possible to assess voltage and time differences, the ratios between the relevant triangle parameters, and the slopes of the lines intersecting its vertices. These measurements contribute to a concrete, multidimensional AP shape description. When the time (*T*_*P*_, *T*_*T HR*_, *T*_*F T R*_) and voltage (*V*_*P*_, *V*_*T HR*_, *V*_*F T RO*_) coordinates are known for all three vertices (P, THR, and FTRO), the necessary information for evaluating these features is available. These properties can be derived directly from raw EP signals obtained in patch-clamp experiments or as precomputed derived properties exposed in EP datasets.

In this work, the focus was on features generated solely from ramp stimulation. Ramps offer the advantage of rapidly and directly generating current-voltage relationships, making them suitable for analyzing cellular responses characterized by rapid activation or time independence [69]. This is particularly relevant for assessing the onset of the AP [69], as ramps help avoid artifacts arising from desensitization [83]. Additionally, the fast exploration of the entire voltage range in voltage-clamp stimulation enables precise threshold detection, and the AP data considered refers to the first AP evoked during ramp stimulation. This is important since, even if the threshold for AP generation exhibits a dynamic range variation depending on the interneuron subtype [40] and the excitability state of the cell, which is regulated through several neuroplasticity mechanisms [18], it ensures the data observed closely represent intrinsic excitability features, preventing the effects of prolonged stimulation, such as adaptation, from introducing confounding factors.

Table 1 presents a set of features that characterize the geometric properties of the triangular NSS representation.

**Table 1.**
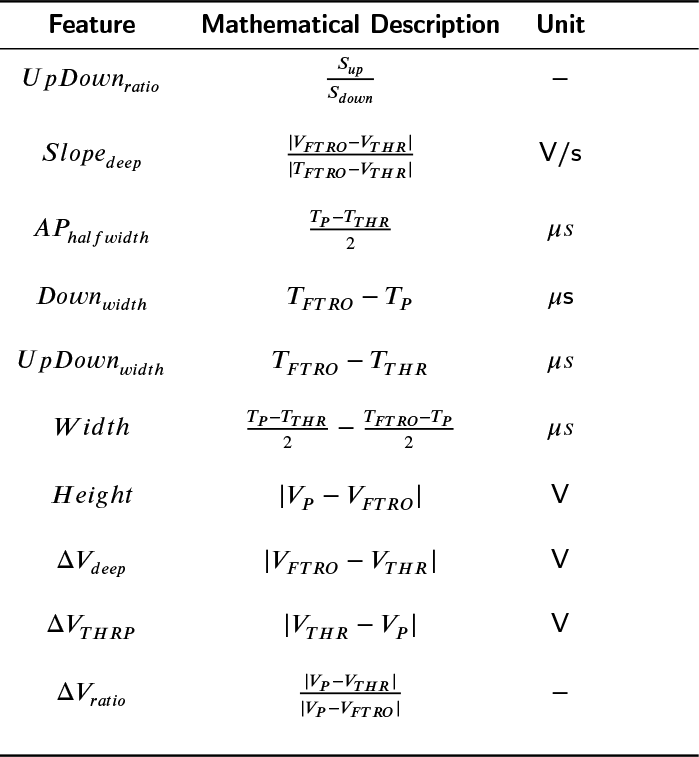
NSS features, along with description and unit of measurement.

NSS features quantitatively express the attributes of the triangular geometric model, with mostly features originating from projections onto the triangle’s axes or direct derivations thereof. Features like *Down*_*width*_ and *U pDown*_*width*_ are projections along the time axis, *Height*, Δ*V*_*deep*_ and Δ*V*_*T HRP*_ represent projections along the voltage axis, while *Slope*_*deep*_, *AP*_*halfwidth*_, *Width* and Δ*V*_*ratio*_ are derived these projections. These operties offer comprehen-sive overview of the depolarization phase (see Figure 1.2), as well as the repolarization and hyperpolarization phases Figure (see1.3-4), providing a synthetic summary of the complete AP shape. It is worth noting that these features focus on modeling the AP shape rather than its position in the time-voltage plane. Therefore, they exclude absolute time and voltage values, emphasizing the relative relationships between absolute voltage and time values at the three key points, expressed as differences or ratios. This differential approach allows for a more generalized description of the AP shape, independent of absolute values. As shown, the NSS framework supports flexible analysis that can adapt to the specific characteristics of the data in different datasets.

Finally, it is important to highlight the bi-univocal mapping between each neuron and the multidimensional point described by the computed NSS features of its AP. This mapping enables the application of clustering and classification algorithms to investigate the heterogeneity of neuronal cells based on their EP profiles alone or in combination with multimodal analysis.

### 4.3. NSS for excitability states identification

Once a cell has been modeled through the EP features constituting NSS, such representation can be used to investigate the existence of cell states across single-cell datasets.

Figure 3 illustrates the flow to perform clustering analysis over NSS features obtained from the datasets. Such exploration is unsupervised, by exploiting clustering approaches to aggregate cells sharing similar AP waveforms, according to NSS representation. Whereas the NSS mapping is pivotal and the innovative core to the provided data exploration pipeline, its analysis relies on state-of-theart clustering techniques. In this case, a well-established method in the literature, i.e., the Hierarchical Agglomerative Clustering (HAC), was selected. This clustering technique begins by treating each data point as an individual cluster. Subsequently, it iteratively merges the closest clusters based on a chosen distance metric. This merging process ultimately results in the creation of a hierarchical structure of clusters referred to as a dendrogram. Strategically cutting this dendrogram to a specific level yields the desired number of clusters for further data analysis. Using HAC offers several advantages: it is not sensitive to initialization conditions (unlike methods like K-Means) or assumptions about cluster shapes; it is robust against outliers in the data; and it allows for examining data groups at different levels of granularity. In this work, Euclidean distance and Ward Linkage are chosen, respectively, as the distance measure to assess the proximity between individual cells and as the merging approach for combining separate clusters during the iterative construction of the final clustering. These two criteria are commonly employed in the literature to identify compact clusters [38].

The choice of clustering cardinality, denoted as *K*, aims to select the number of clusters that better supports data exploration. While with HAC it usually involves analyzing the dendrogram generated by the algorithm, in this pipeline *a priori* optimization using maximization of the Silhouette score [65] is selected to standardize the procedure. The Silhouette metric *s* is calculated as follows:

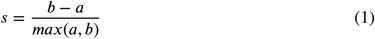

where the average intra-cluster distance *a* quantifies intra-cluster cohesion, and the average inter-cluster distance *b* measures inter-cluster separation. Unlike other metrics such as the Sum of Square Errors (SSE), the Silhouette metric, given its mathematical definition, is robust to nonspherical clusters [38].

### 4.4. Validation of NSS clusters as excitability states

A multi-step validation shows that the proposed methodology identifies intrinsic excitability states, targeting the following aspects:

#### 1. Coherence with excitability definitions

Analysis of the distribution of cells within each NSS cluster over features that are traditionally used in the literature to model excitability: (i) Action Potential Half Width (AP_half−width_), i.e., the duration of the initial part of the AP in ms; (ii) Inter Spike Interval (ISI), the average time interval separating APs in the cell’s firing response in ms; (iii) f-I curve slope (FICS), the slope of the f-I curve, putting the firing rate in relation to stimulation current intensity; and (iv) *I*_*T HR*_, i.e., the minimum current intensity level capable of generating a firing response, in pA.

#### 2. Coherence with known excitability states

Analysis of the prevalence of cell-type labels within each NSS cluster to map the relationships between NSS clusters and cell types with recognized intrinsic excitability properties. Cell-type labels are provided as metadata in the datasets.

#### 3. Characterization of excitability states

Investigation of the relevance of NSS features to explain clustering results in relation to intrinsic excitability states. This analysis relies on two approaches:

- Spearman’s correlation [66], a statistical method to measure the strength and direction of the *linear* relationships between the NSS features and the identified intrinsic excitability states. The Spearman’s correlation coefficient yields a numerical value ranging from -1 to 1, where a positive value indicates a positive correlation and a negative value indicates a negative correlation.
- Permutation feature importance method based on an Random Forest (RF) classifier [7] to detect *non-linear* relationships between the NSS features and the identified intrinsic excitability states. RF is a versatile machine learning algorithm capable of assessing feature importance within a dataset [64]. In essence, it evaluates how each feature contributes to prediction accuracy across an ensemble of decision trees. This approach offers a robust and interpretable means of identifying the most pertinent features.

### 4.5 Transcriptional exploration of NSS clusters

While the main NSS validation focuses on the analysis of EP features, further insights may be collected looking at the transcriptional profiles of cells belonging to the NSS clusters. This is particularly true when analyzing new unknown states identified by NSS.

The transcriptional exploration employs the transcriptomic profiles of cells to explore the NSS clustering results. This specific analysis is applicable to Patch-seq datasets, which provide electrophysiological and gene expression data simultaneously on the same cells (see Section 2), facilitating cross-validation and joint analysis of EP and transcriptomic findings. The analysis follows a workflow for scRNA-seq data analysis that is well-established in the research community [44], using the Seurat pipeline [27] to cover the following steps:

#### 1. Dimensional reduction

This step generates a twodimensional (2D) visualization of the embedding of the cells generated with Uniform Manifold Approximation and Projection (UMAP), a non-linear dimensional reduction method highlighting the global differences of the transcriptomic profiles. All the following steps base upon this transcriptomic embedding.

#### 2. Cell types identification

This step links cells to the four distinct GABAergic neuronal subtypes by labeling cells with expression levels of cell-type marker genes and cell-type labels, respectively. These four are very well known and characterized subtypes of cortical GABAergic neurons, Pvalb, Sst, Vip, and Lamp4. Their identification supports the subsequent analyses and exploration of the NSS clusters by supporting their identity as excitability states. Indeed, as discussed in section 3, the Pvalb cell type has a typical pattern of higher excitability (higher spiking rates, yet higher threshold) with respect to the other subtypes, thus its prevalence in a NSS cluster supports its representation power of excitability. The *PatchClampDataset* provides gene expression levels as transcriptomic data and cell type labels as metadata [24].

#### 3. NSS clusters over cell types comparison

The following step examines the distribution of NSS labels across transcriptomic cell types and subtypes by comparing their respective visualizations on the transcriptomic embedding.

#### 4. Intra-type NSS clusters Differential Expression (DE) analysis

Transcriptomic data allow DE analysis among NSS clusters to investigate gene expression profiles that could shed light on the underlying biological mechanisms for each identified state. This can be applied at different clustering resolutions. Specifically, within the same transcriptomic subset, DE analysis targets groups of cells with different NSS labels to identify intra-type differences. The analysis provides a list of the top DE genes, highlighting the genes that characterize each group primarily and specifically compared to the other.

#### 5. Ontology-based biological processes enrichment

Gene Ontology (GO) Enrichment Analysis (EA) [52] supports the further and systematic investigation of the resulting lists of DE genes. This approach helps to generate hypotheses based on knowledge about the involvement of specific gene sets in biological processes. The GO is a systematic and curated nomenclature and annotation of biological processes, molecular functions, and cellular components associated with each gene. The GO EA of a gene list highlights overrepresented processes and functions in GO. Additionally, the provided analysis involves a manually curated investigation of the functional aspects of specific genes. This work uses the GO platform supported by PANTHER [52], using the ShinyGO tool [21] to perform systematic GO EA, and leverages on GeneCards [70] or Uniprot [15] platforms for the subsequent investigations.

### 4.6 Datasets

The presented analysis relies on two datasets of murine visual cortical neurons. Table 2 illustrates their characteristics. The scarcity of datasets containing a sufficient cell count is a notable challenge within the realm of patch-seq data. The provided study exclusively centers on murine cellular data, which were thoughtfully selected based on criteria such as cell count, data quality, and accessibility.

**Table 2.**
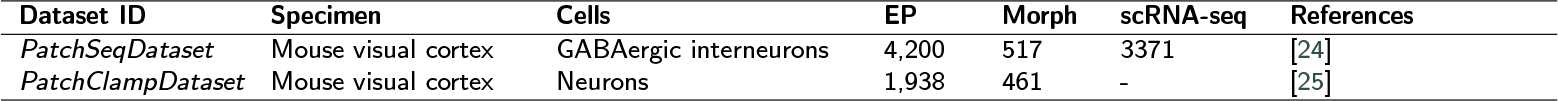
Datasets for NSS analysis. For each dataset, the number of cells profiled under EP, morphological and transcriptomic profiles is provided. EP: Electrophysiological profile; Morph: morphological reconstructions; scRNA-seq: transcriptomic profile.

The *PatchSeqDataset* [24] takes advantage of the patchseq technique, which combines patch-clamp with scRNA-seq to collect EP and transcriptomic profiles of 4,200 mouse visual cortical GABAergic interneurons, reconstructing the morphological conformation of 517 of them.

The *PatchClampDataset* [25] provides EP data from 1,938 neurons of adult mouse visual cortical neurons and morphological reconstructions for 461 of them. This dataset, accessible online via the *Allen Brain Atlas Cell Types Database* [2], provides a mixture of spiny and aspiny neurons. This work analyzed only cells labeled as *aspiny*, under the consensus that this morphological label is a strong indication that the cell is an interneuron.

The experiments proposed in this paper focus on the GABAergic neurons from both datasets. Notably, GABAergic neurons are abundantly represented in the *PatchSeqDataset* [24], whereas they comprise approximately half of the cell population in the *PatchClampDataset* [25]. The selection of GABAergic neurons resorts to the metadata provided in the datasets: transcriptomic cell type labels supplied by the authors allowed the consideration of only cells labeled as GABAergic neurons for *PatchSeqDataset* [24], and morphological metadata allowed the consideration of only cells labeled as *aspiny* (thus, putatively GABAergic) for *PatchClampDataset* [25], respectively.

As a premise for the provided analysis, a data cleaning step removed cells with incomplete information or outlier feature values in both datasets. Taking into account the cleaning and filtering of data that retain only GABAergic cells, the analysis employs 3,653 neurons for *PatchSeqDataset* and 770 for *PatchClampDataset*.

The main analysis presented in this study focuses on the *PatchSeqDataset*, as it provides both EP and transcriptomic data, enabling direct cross-validation of the NSS-based excitability states identification with transcriptomic ground truths. The *PatchClampDataset* supports the further EP validation of the NSS method, but, since it only provides coarse-grained labeling of cell types using transgenic murine lines (see Section 5), it does not support transcriptomic validation.

Both *PatchClampDataset* and *PatchSeqDataset* provide comprehensive EP characterization of single cells, including voltage-clamp stimuli and responses collected for each sample. Each cell in the datasets is associated with multiple types of stimulation, corresponding to standard patchclamp protocols such as current ramps, short squares, and long squares. For each stimulation type, there are multiple *sweeps*, which are repeated trials in which the parameters of the stimulation protocol vary, resulting in different EP responses. For example, the long square stimulation set includes a sweep for each step of the current amplitude applied as input to the neuron.

## 5. Results and Discussion

This section proposes experimental results from the application of NSS on both the *PatchSeqDataset* and the *PatchClampDataset*. While the *PatchSeqDataset* supports full multimodal analysis, the *PatchClampDataset* only provides EP analysis. Results validate NSS’s capability for identifying both the well-known fast-spiking state and potential new candidate excitability states within both the considered datasets. New identified states are thoroughly characterized based on their electrophysiological and transcriptomic profiles.

### 5.1 Validation: NSS identifies the fast-spiking excitability state

NSS was validated by proving its capability of identifying well-known excitability states in both datasets and identifying new, unknown ones independently of cell types.

Following the approach described in Section 4 (see Figure 3), HAC was applied to computed and z-score standardized NSS features from Table 1 on both datasets. For *PatchSeqDataset*, the optimization of the clustering identified an optimal number of *K* = 3 clusters (Silhouette score: 0.34). A second solution, *K* = 2, achieved a similar score (Silhouette score: 0.32) but was neglected in favor of the best-performing cardinality. For *PatchClampDataset*, the clustering analysis produced two potential solutions: the optimal clustering was identified at *K* = 2 (Silhouette=0.40), slightly better than *K* = 3 (Silhouette=0.38).

Figure 4 illustrates the scatter plots of the data based on the Principal Component Analysis (PCA) analysis of NSS features for *PatchSeqDataset* (top left panel) and *PatchClampDataset* (top right panel), respectively. The dot colors refer to the identified clusters. Clustering results for *PatchSeqDataset* (*K* = 3) denote clusters as *EC0* (violet), *EC1* (green), and *EC2* (yellow). Clustering results for *PatchClampDataset* (*K* = 2) denote clusters as *EC0* (violet) and *EC1* (yellow).

Considering that GABAergic neurons, the main focus of this analysis, present four cell subtypes, to exclude the possibility that in the *PatchSeqDataset* NSS was merely recapitulating transcriptomic clustering, the *K* = 4 clustering case was also analyzed. However, this case only introduced a further split within the *EC1* cluster. This additional grouping neither recapitulates transcriptomic cell-type regions nor provides more informative insights than the other case (data not shown).

State-of-the-art methods to identify the fast-spiking state resort to the excitability features introduced in Section 4.4. In particular, the fast-spiking state is characterized by low AP_half−width_ and ISI, high FICS, and high *I*_*T HR*_. Through a comparative analysis of the fast-spiking feature values across clusters, the remaining rows in Figure 4 show that

NSS captures the fast-spiking excitability state and additional unknown states. In particular, cluster *EC0* captures the fast-spiking state in both datasets, consistently exhibiting the lowest AP_half−width_ and ISI, the highest FICS, and the highest *I*_*T HR*_ compared to the other clusters within the same dataset. Additionally, NSS captures other unknown states in both *PatchSeqDataset* (clusters *EC1* and *EC2*) and *PatchClampDataset* (cluster *EC1*), further discussed later.

At this stage, it is important to confirm that NSS actually identifies excitability states, not cell types. This passes through the assessment that the NSS cluster *EC0* captures the fast-spiking state, not just a single cell type. The fastspiking state and the Pvalb interneuronal cell subtype have a strong link. Among GABAergic interneurons, the Pvalb subtype is associated with the fast-spiking excitability state. Fast-spiking cells have intense activation upon stimulation and high thresholds. This unique functional profile appears to support the role of Pvalb cells as primary regulators of excitable neurons, as highlighted in [13]. This role demands that they fire with high intensity to exert an effect (high activation) while selectively integrating stimuli from other GABAergic interneurons in the inhibitory network (high threshold).

Figure 5 overlays the *I*_*THR*_ distributions of the different GABAergic subtypes onto the scatter plots of cells with NSS cluster labels for the *PatchSeqDataset*. The identification of the cell types relies on the gene expression profiles of cells. The results show that almost all Pvalb cells have an *EC0* label. Since Pvalb cells are notably in the fast-spiking state, this finding supports that the NSS cluster *EC0* corresponds to the fast-spiking excitability state. Nevertheless, cluster *EC0* is not limited to a single cell type. Sst cells also significantly contribute to the definition of this cluster through cells with high *I*_*T HR*_ values (i.e., high *I*_*T HR*_ is a marker of the fast-spiking state) further confirming that *EC0* represents a state and not a cell type. These findings are consistent with results obtained in human cells, where the electrophysiological profile of some Sst cells shows they are in a fast-spiking state, drawing a close similarity to Pvalb cells [39]. In conclusion, results validate NSS capability of capturing the well-known fast-spiking excitability state independently of cell types.

**Figure 5:**
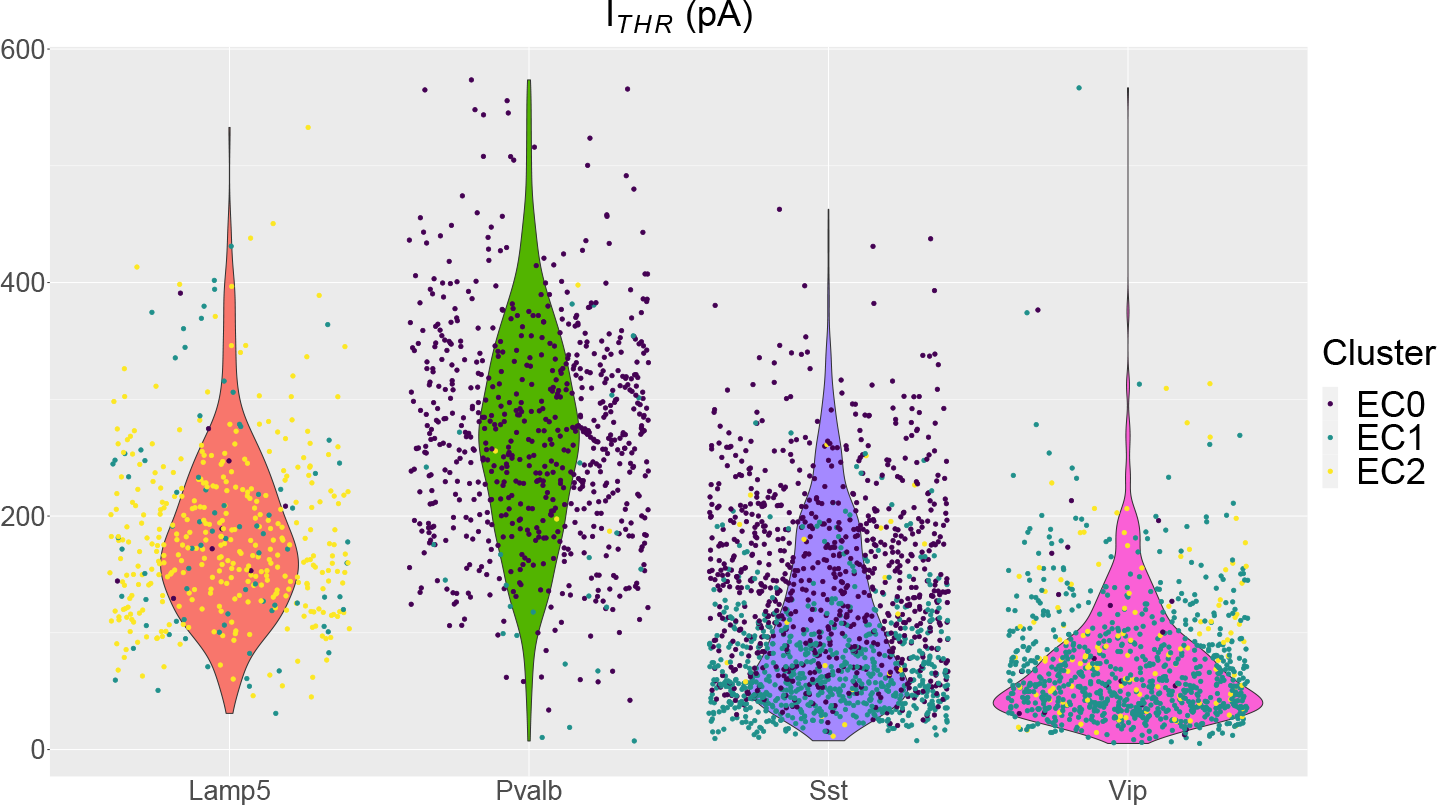
*I*_*T HR*_ distribution and NSS clustering labels across GABAergic subtypes. Violin plots show the distribution of cells within each subtype over *I*_*T HR*_ feature values, scattered dots are cells over such distributions, and their colors represent NSS cluster labels. The color coding refers to clustering labels: violet for *EC0*, green for *EC1* and yellow for *EC2*. Cell-type labels are based on *PatchSeqDataset* metadata. Cells labeled as “Serpinf1”, “Sncg” and “Meis2” are grouped under the broader cell subtype “Vip”.

A similar analysis performed on the *PatchClampDataset* produced coherent results. 71% of *EC0* cells are labeled as Pvalb cells, and 10% are Sst cells with high *I*_*T HR*_. The Sst cells are the 22% of *EC1* cells, showing that they belong to both *EC0* and *EC1* clusters. Since the *PatchClampDataset* does not contain transcriptomic data, cell types were assigned using labels from Cre lines of the transgenic mice used in the patch-clamp experiments [25].

To further exclude that NSS is identifying cell types, Figure 6 shows the embedding of the Sst cells labeling the cells with subtypes obtained from metadata available in the *PatchSeqDataset* [24] (left side). On the right side, the same embedding is labeled based on the NSS clusters. Again, the figure confirms that cluster *EC0* is not associated with a specific cell subtype.

**Figure 6:**
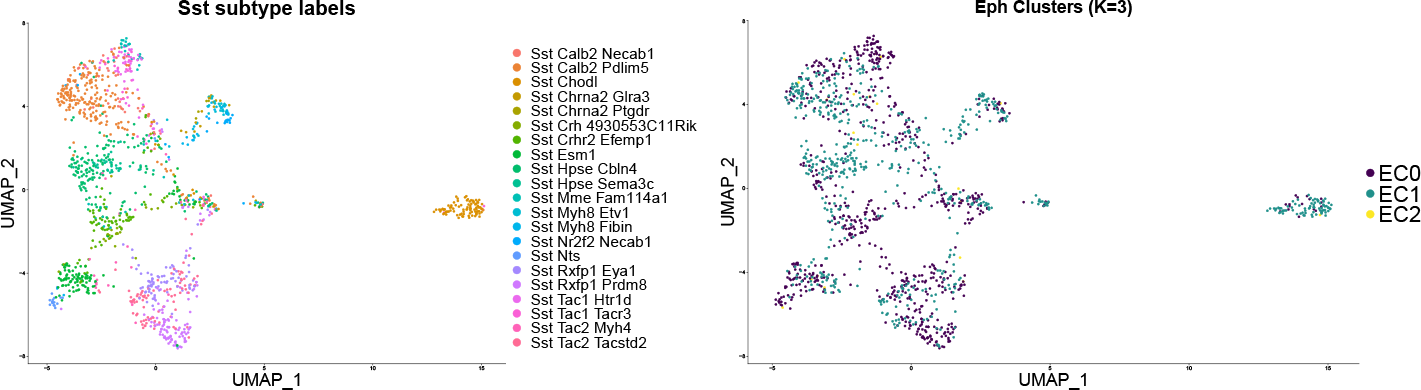
Sst cells labeled with the transcriptional cell sub-types provided by the metadata (on the left) and the EP clusters (on the right).

An additional validation of the correct identification of the fast-spiking state among the Sst cells can be obtained through Gene Ontology (GO) Enrichment Analysis (EA).In fact, the fast-spiking state presents positive regulation of processes involving potassium (*K*^+^) channels. They sustain the intense firing activity typical of the fast-spiking state by shortening the AP repolarization and hyperpolarization phases, that is, the refractory period, where AP firing is inhibited [45].

A GO EA performed over the 111 positively DE genes in *EC0* versus *EC1* cells within the Sst subset highlighted that, within the Sst embedding, the *EC0* cluster exhibits an enrichment of *K*^+^ channels-related processes, consistent with the fast-spiking state it captures. In particular, Figure 7 shows the enrichment of specific ontology elements related to *K*^+^ channels, particularly in the *molecular functions* ontology, where “Voltage-gated *K*^+^” and “*K*^+^ channel activity” are enriched. The DE genes list contains numerous entries associated with *K*^+^ channels, which are characterized by the “*Kcn*” naming pattern [67]. This further validates that *EC0* captures the fast-spiking state. Since the comparison is based on Sst cells only, results pertain to the fast-spiking state and not to cell subtypes.

**Figure 7:**
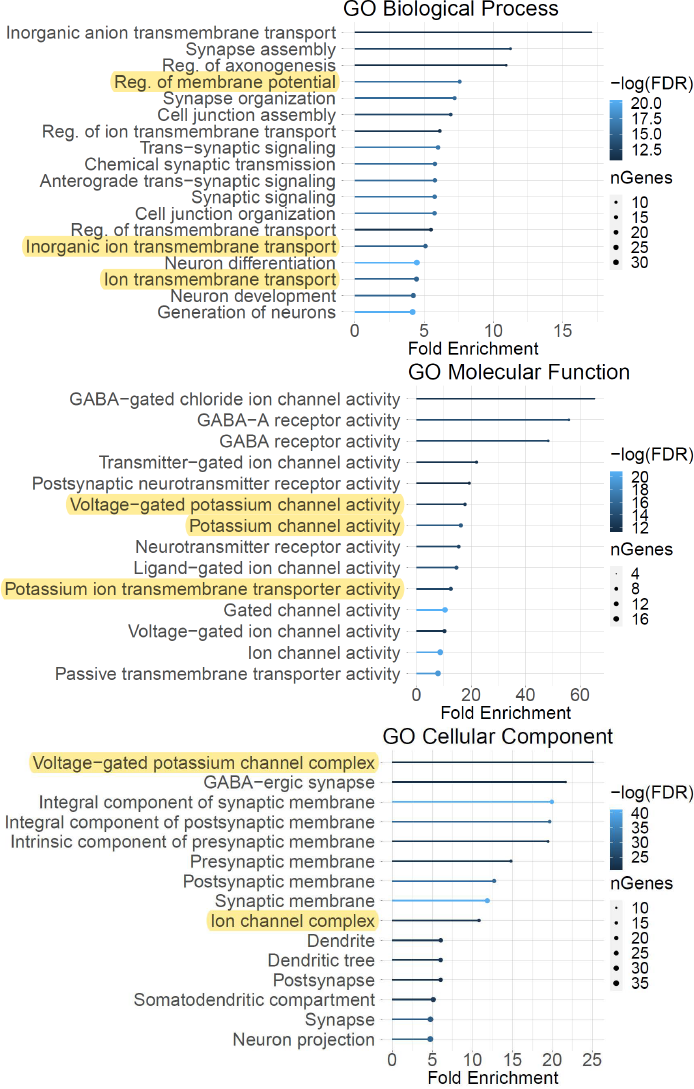
GO EA analysis of the *EC0* versus the *EC1* cluster within the Sst cells subset. Results are provided for Biological Process (Top), Molecular Function (Center), and Cellular Component (Bottom). The *x* axis represents the fold change, ranked from the highest. The highlighted terms are the ones directly correlated to *K*^+^ channels.

### 5.2 Exploration: NSS supports the characterization of known and unknown excitability states

The validations conducted so far have affirmed the capability of NSS in not only identifying the well-known fastspiking state but also in discerning new states, denoted as clusters *EC1* and *EC2* for *PatchSeqDataset*, and cluster *EC1* for *PatchClampDataset*. This section performs further investigations to understand if these novel states are potential candidates for new excitability states.

Distinctive attributes in the repolarization and hyperpolarization phases of the Action Potential, alongside the gene expression patterns of potassium channels, notably contribute to the regulation of various neuronal excitability states. States identified by NSS primarily diverge in their AP repolarization and hyperpolarization phases, displaying varying expression profiles of potassium channels that modulate the duration of the refractory period. This aspect strongly substantiates their classification as novel neuronal excitability states.

To support this investigation, Figures 8 and 9 visually represent the Spearman’s linear correlation of NSS features with the cluster label (column *LABEL*) for the *PatchSeqDataset* and the *PatchClampDataset*, respectively. Features such as Δ*V*_*deep*_, Δ*V*_*ratio*_, and Δ*V*_*thrp*_, associated with the AP repolarization and hyperpolarization phases, exhibit the highest linear correlation with cluster labels, suggesting that the information encapsulated within these elements explicates a substantial portion of the inter-cluster variance. Given the pivotal role played by these phases in characterizing neuronal excitability, this finding effectively supports that the identified clusters capture distinct excitability states.

**Figure 8:**
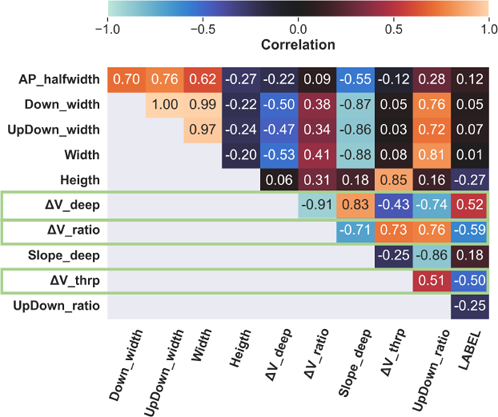
Spearman’s correlation of NSS features and cluster labels for PatchSeqDataset. An hot (high positive correlation) and cold (high negative correlation) diverging color map represent the correlation levels.

**Figure 9:**
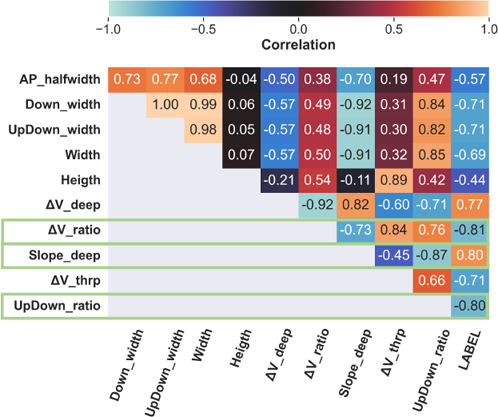
Spearman’s correlation of NSS features to clustering label-excitability state. An hot (high positive correlation) and cold (high negative correlation) diverging color map represent the correlation levels.

Figures 10 and 11 further support this finding reporting the mean accuracy decrease of RF classification upon the removal of specific features from the model for the *PatchSeqDataset* and the *PatchClampDataset*, respectively. Notably, the removal of the Δ*V*_*ratio*_ led to an approximate 4% decrease in mean accuracy for the *PatchSeqDataset* and an 11% decrease for the *PatchClampDataset*. This trend reinforces the criticality of features associated with the AP repolarization and hyperpolarization phases, solidifying the NSS clusters as indicative of distinct neuronal excitability states. Indeed, varying durations of AP repolarization directly influence the refractory period, thus correlating with the regulation of diverse excitability states [45].

**Figure 10:**
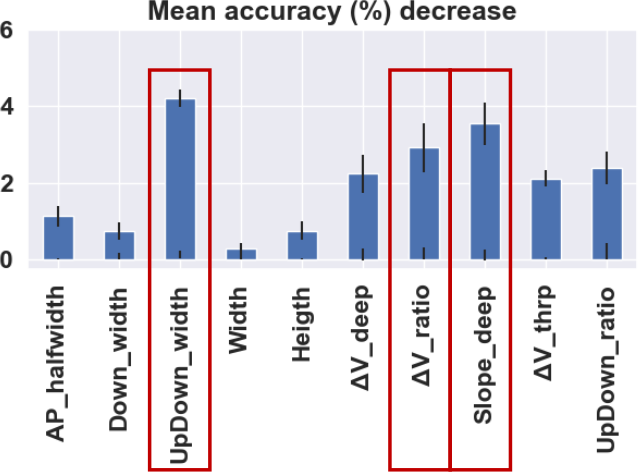
Mean accuracy decrease in the cluster label prediction for *PatchSeqDataset* done by RF model when shuffling values of NSS features

**Figure 11:**
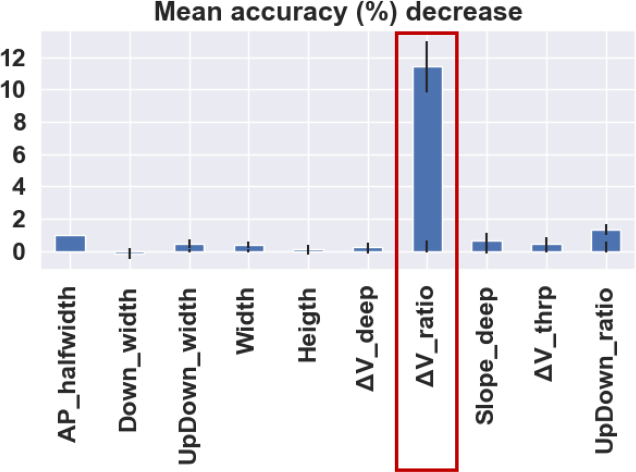
Mean accuracy decrease in the cluster label prediction for *PatchClampDataset* done by RF model when shuffling values of NSS features.

Consistent with the results of the feature importance analysis and their role in excitability regulation, the presence of different excitability states was also analyzed by comparing the typical AP waveform of each NSS cluster, searching for significant differences in the repolarization and hyperpolarization phases.

Figures 12 and 13 display the raw AP waveforms of the median cells within each cluster (the closest cells to the centroid of the respective cluster) for *PatchSeqDataset* and *PatchClampDataset*, respectively.

**Figure 12:**
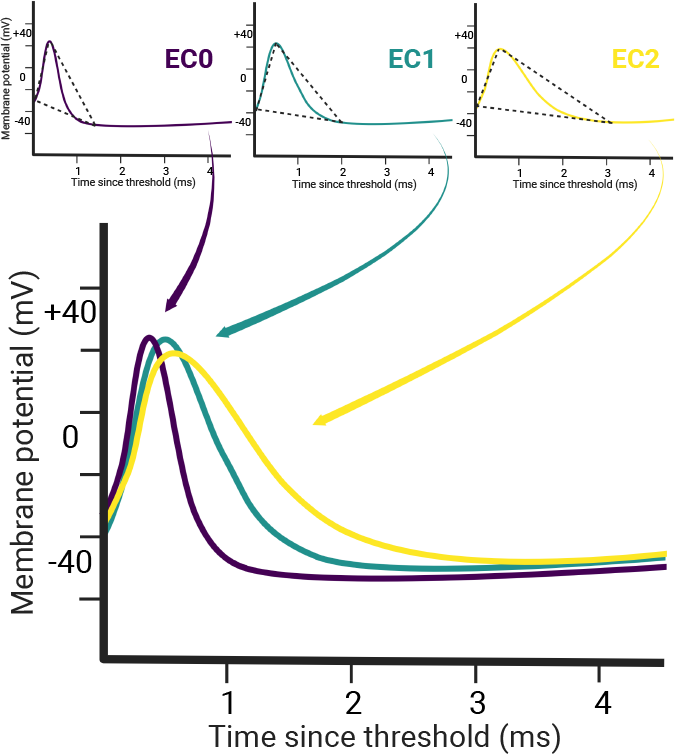
AP spikes of median cells identified using NSS on *PatchClampDataset* with coloring with respect to label of *K* = 2 NSS clustering: *EC0* (violet), *EC1* (green),*EC2* (yellow).Median AP spikes and their superimposition. NSS backbones THR-P-FTRO triangles are reported as dashed line over the spiking graphs.

**Figure 13:**
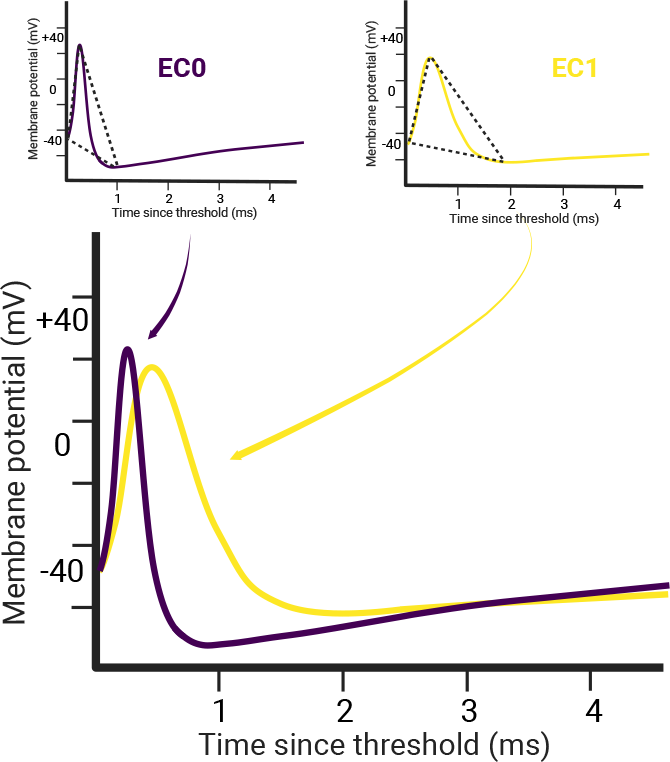
AP spikes of median cells identified using NSS on *PatchClampDataset* with coloring with respect to label of *K* = 2 NSS clustering: *EC0* (violet), *EC1* (yellow).

For the *PatchSeqDataset*, the comparison of the AP waveforms from clusters *EC0, EC1*, and *EC2* (Figure 12) reveals three distinct AP shapes, where the repolarization and hyperpolarization phases exhibit the most variability across clusters, particularly in the region near the FTRO, corresponding to a different typical duration for the refractory period. The NSS *EC0* cluster displays a narrow AP shape with a deep FTRO, while the *EC2* cluster demonstrates the opposite behavior, characterized by a gradual repolarization phase and an almost flat FTRO. Finally, the *EC1* cluster occupies an intermediate position between these two extremes.

For the *PatchClampDataset* (Figure 13), the AP shape of the median cells from *EC0* demonstrates a rapid decline in the repolarization phase, while the AP shape from *EC1* exhibits the opposite trend, with a longer AP duration and a slower repolarization phase, corresponding to a longer refractory period.

This result confirms the relevance of the repolarization and hyperpolarization phases in distinguishing the NSS clusters, supporting their classification as distinct excitability states.

Eventually, the differences in the gene expression patterns of potassium channels significantly influence the repolarization and hyperpolarization phases of the AP, emphasizing distinct neuronal excitability states. The NSS clusters showcase three distinct gene expression patterns for potassium channels, solidifying the identification of NSS clusters as excitability states at both the electrophysiological and molecular levels.

Figure 14.A and Figure 14.B illustrate the gene expression profiles of two *K*^+^ channel-related genes (i.e., *Kcnc2* and *Kcnn2*) for the three clusters of the *PatchSeqDataset*. Both genes exhibit expression levels that support *EC0* as the fast-spiking excitability state, as well as *EC1* and *EC2* as distinct excitability states. *Kcnc2*, associated with fast AP repolarization, demonstrates the highest expression levels in *EC0*, intermediate levels in *EC1*, and the lowest levels in *EC2* (Figura 14.A). This observation confirms *EC0* as the fast-spiking state, where increased expression of specific *K*^+^ channel genes indicates a positive regulation of AP repolarization steepness, resulting in a narrower AP shape. Likewise, it validates *EC1* and *EC2* as intermediate and lowest excitability states, respectively.

**Figure 14:**
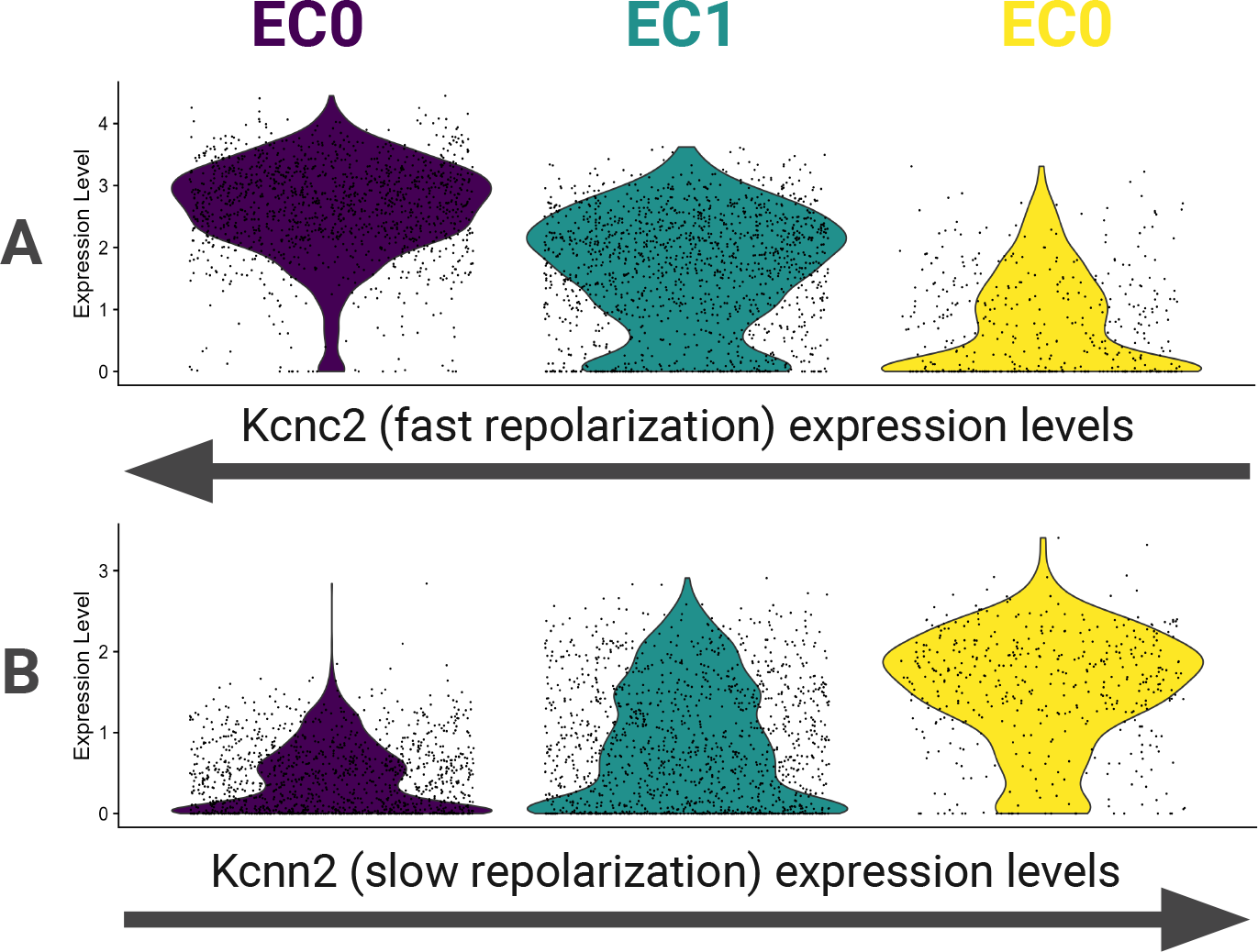
Relations between the median AP shapes with exemplary triangles (first row), the NSS clustering labels over the transcriptomic embedding, with an indication of the transcriptional GABAergic subtypes associated to each cluster (second row), and the expression levels of exemplary *K*^+^-channels-related genes (*Kcnc2* and *Kcnn2*) across NSS clusters, represented through violin plots showing the gene expression levels distributions across cells in each cluster. The colors identify the NSS clusters for the *K* = 3 case, namely violet for *EC0*, green for *EC1*, and yellow for *EC2*.

Conversely, *Kcnn2* contributes to the slow component of the synaptic after-hyperpolarization phase. As a member of the SKCa channels, *Kcnn2* serves as a vital mediator of spike adaptation by generating a substantial afterhyperpolarization in instances of intense activity [26]. This phase primarily involves *Ca*^++^-dependent and voltage-independent *K*^+^ currents that can persist for several seconds following a series of APs. Thus, the observed reverse trend in the expression levels of *Kcnn2* indicates the positive regulation of a slower repolarization phase in *EC2* and *EC1*. This once again confirms that *EC0* represents the fast-spiking state, which corresponds to the absence of firing adaptation.

Taking together all these findings NSS identifies three excitability states in *PatchSeqDataset. EC0* cluster, as extensively validated in Section 5.1, captures the fast-spiking excitability state; *EC2* captures a low excitability state and *EC1* captures an intermediate state between the two.

## 6. Conclusions

This paper introduced the NSS approach, which offers a straightforward method to analyze AP waveforms for studying neuronal cell states. The article presents the framework of NSS analysis, outlines its data requirements and demonstrates its application using two data sets of murine neocortical neurons *PatchClampDataset* and *PatchSeqDataset*, described in [25] and [24], respectively. The NSS analysis captures cellular heterogeneity and in particular cell-type independent neuronal excitability states. In fact, it identifies a sub-portion of Sst cells in a fast-spiking excitability state, consistently to what observed in [39] for human cells. NSS partitions prove consistent with existing activity-based, and not threshold-based definitions of neuronal excitability. In particular, in both datasets, NSS identifies a cluster that captures the well-characterized fast-spiking neuronal excitability state that, in *PatchSeqDataset*, proves to be specifically enriched for cellular processes related to *K*^+^ channel metabolism and plasticity. This is consistent with the notion that *K*^+^ channels play a crucial role in regulating AP waveforms, particularly in the repolarization and hyperpolarization phases, which explain most of the variability observed across the identified clusters. The EP analysis combined with cell line-based cell type labeling over *PatchClampDataset* provides additional validation of the NSS capability of capturing the heterogeneity of cells with respect to types and states that are consistent across the two datasets.

NSS facilitates EP analysis and can be applied to singlecell patch-clamp datasets. Patch-seq datasets also enable multimodal analysis and biological validation of the NSS approach. The NSS-based analysis benefits from patchseq datasets due to their inherent consistency between EP and transcriptomic cellular profiles. Unfortunately, patchseq datasets currently face challenges in terms of data availability, and they often have low throughput due to technical limitations, resulting in a limited number of samples per dataset and hindering large-scale analysis [42, 48]. Nevertheless, recent advancements in semi-automatic patchclamp setups allow for the generation and publication of large-scale patch-clamp datasets [49].

Apart from the limitations inherent to patch-seq data, which include issues like data scarcity (where only a handful of datasets, around 3-4, support multimodal analysis) and challenges related to the accessibility and interoperability of available patch-seq datasets, NSS also suffers from limitations connected to the simplicity of the model, that on the other had is also one of the strengths. It is important to note that NSS focuses on just one of the numerous electrophysiological processes within cells. While this approach yields simple and comprehensible results, it provides a limited perspective on neuronal complexity. Nonetheless, considering this as the initial step in a modular approach to dissecting neuronal complexity, we anticipate the development of complementary components to NSS that will broaden the coverage of our overall strategy. Limitation of this work also include the narrow manually curated analysis of transcriptomic results, which focuses on the most relevant findings.

Thus, the forthcoming advancements encompass the application of the method to new patch-seq datasets as they become accessible, the expansion of the analysis of neuron dynamics with the incorporation of new modular components, and the enhancement of the multimodal validation process and manually curated analysis to make it more systematic and comprehensive.

Additionally, once comprehensive human patch-seq datasets become available on a larger scale, we plan to extend the validation of the NSS method to include them. In particular, we aim to apply the NSS methodology on the patch-seq dataset analyzed in [39], where the electrophysiological profile of one of the Sst subtypes exhibits similarity to the Pvalb profile, that is, to a fast-spiking excitability state as soon as it will be made publicly available. Future research aims to analyze more datasets containing different neuronal types and potentially from other species. This includes both patch-clamp and patch-seq datasets. Future research will incorporate a more comprehensive characterization of gene expression profiles within NSS-based partitions regarding multimodal analysis. Moreover, the future availability for human neuronal data, will provide ground for testing the NSS method not only on murine dataset, showing its generality. This will allow for the detection of complex expression patterns and expand subsequent analysis to a broader range of genes. Such an approach will provide a more extensive exploration of the NSS biological significance, including considering alternative or integrative approaches to ontology-based analyses.

Many neuronal properties and behaviors lack a universally agreed-upon definition and comprehensive characterization in the existing literature. The NSS approach can contribute to the analytical study of these properties within patch-seq datasets, helping to establish consensus around essential and widely used concepts in neuronal physiology. An important future direction is to utilize the NSS approach to investigate the relationships between AP shape and crucial neuronal features beyond the sole intrinsic excitability, as provided in this work, and analysing also other key aspects such as *plasticity* [18] and *firing rate set points* [76] in neuronal cells.

## Code availability statement

Electrophysiological analysis and NSS-based clustering based on the Python 3 programming language, multimodal validation combines Python 3 and R programming languages. The code supporting this study is publicly available on GitHub: https://github.com/smilies-polito/NSS.

### No competing interests statement

The authors declare that they have no known competing financial interests or personal relationships that could have appeared to influence the work reported in this paper.

## CRediT authorship contribution statement

**Lorenzo Martini:** Conceptualization, Methodology, Software, Validation, Formal Analysis, Investigation, Writing - Original Draft, Writing - Review & Editing, Visualization. **Gianluca Amprimo:** Conceptualization, Methodology, Software, Validation, Formal Analysis, Investigation, Writing - Original Draft, Writing - Review & Editing, Visualization. **Stefano Di Carlo:** Conceptualization, Methodology, Resources, Writing - Review & Editing, Supervision, Funding acquisition. **Gabriella Olmo:** Conceptualization, Methodology, Resources, Writing - Review & Editing, Supervision, Funding acquisition. **Claudia Ferraris:** Conceptualization, Methodology, Resources, Writing - Review & Editing, Supervision, Funding acquisition. **Alessandro Savino:** Methodology, Resources, Supervision, Funding acquisition. **Roberta Bardini:** Conceptualization, Methodology, Writing - Original Draft, Writing - Review & Editing, Supervision, Project administration.

---

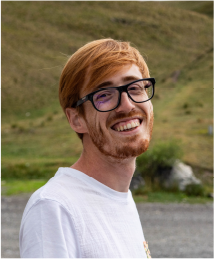

Lorenzo Martini is currently a Ph.D student at the Control and Computer Engineering Departement of politecnico di Torino. He holds a M.Sc. in Physiscs of COmplex Systems from Politecnico di Torino. His research centers on bioinformatics and computational biology, with a current focus on single-cell genomics and multimodal analysis.

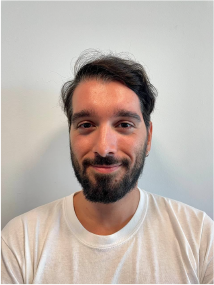

Gianluca Amprimo received his Master’s Degree in Computer Engineering from the Politecnico di Torino in 2020. He is currently a PhD student at the Control and Computer Engineering Department of Politecnico di Torino and a Research Fellow at the National Research Council (CNR-IEIIT) of Italy. His main research interests include human pose estimation from video, innovative technologies for telemonitoring and rehabilitation, and AI for medical application.

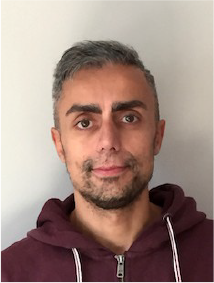

Stefano Di Carlo is a full professor in the Control and Computer Engineering department at Politecnico di Torino (Italy) since 2021. He holds a Ph.D. (2003) and an M.S. equivalent (1999) in Computer Engineering and Information Technology from the Politecnico di Torino in Italy. Di Carlo’s research contributions include biological network analysis and simulation, machine learning, image processing, and evolutionary algorithms, as well as development of resilient computer architectures.

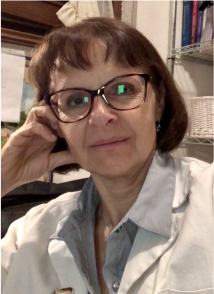

Gabriella Olmo (IEEE Senior Member) received her M.E. and Ph.D. degrees in Electronic Engineering from Politecnico di Torino, Italy, in 1986 and 1992, respectively. In 2016, she received her Master’s degree in Medicine and Surgery from Universit’a di Torino, Italy. She is currently a full professor in the Department of Control and Computer Engineering, Politecnico di Torino, Italy. Her main research interests are in the fields of wearable sensors, signal processing, and machine learning techniques for medical applications. She is coauthor of more than 250 publications in international journals and proceedings in international conferences.

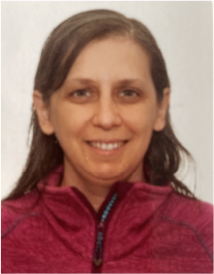

Claudia Ferraris graduated (summa cum laude) in Computer Science from the University of Turin (Italy) in 1997. After graduation, her first activities were at CNR research centers, working on image/video coding, compression techniques and motion estimation algorithms. After several years in the aeronautical and automotive industries, she was a research fellow at the Institute of Electronics, Information Engineering and Telecommunications (IEIIT) of CNR from 2012 to 2020, where she began working on health and well-being, noninvasive technologies for motion analysis, remote monitoring, and rehabilitation in pathological conditions. She has been a permanent researcher at the same institute since 2020 and has expanded her expertise, focusing on exergaming, body tracking, RGB-Depth sensors, artificial intelligence, machine learning, and deep learning methods. She received her PhD in Neuroscience from the University of Turin (Italy) in 2021. She has authored numerous research papers published in international conference proceedings, national and international journals.

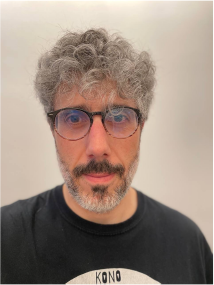

Alessandro Savino is an Associate Professor in the Department of Control and Computer Engineering at Politecnico di Torino (Italy). He holds a Ph.D. (2009) and an M.S. equivalent (2005) in Computer Engineering and Information Technology from the Politecnico di Torino in Italy. Prof. Savino’s research contributions include Approximate Computing, Reliability Analysis, Safety-Critical Systems, Software-Based Self-Test, and Image Analysis. He has been part of the program and organizing committee of several IEEE and INSTICC conferences and served as a reviewer of IEEE conferences and journals. His research interests include Operating Systems, Imaging algorithms, Machine Learning, Evolutionary Algorithms, Graphical User Interface experience, and Audio manipulation.

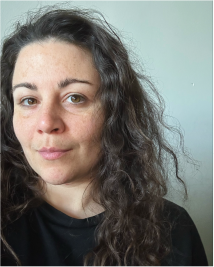

Roberta Bardini is a post-doc researcher at the Department of Control and Computer Engineering of Politecnico di Torino. She holds a Ph.D. (2019) in Control and Computer Engineering from the Politecnico di Torino in Italy, and a M.Sc. in Molecular Biotechnology (2014) from Università di Torino. Her research focuses on computational biology and bioinformatic approaches for analyzing and modeling complex biological systems, and on the computational design and optimization of biomanufacturing processes.

## References

[1] Arlotta, P., Paşca, S.P., 2019. Cell diversity in the human cerebral cortex: from the embryo to brain organoids. Current opinion in neurobiology 56, 194–198.

[2] Atlas, A.B., 2023. Cell types database. URL: https://celltypes.brain-map.org/.

[3] Barnett, M.W., Larkman, P.M., 2007. The action potential. Practical neurology 7, 192–197.

[4] Bean, B.P., 2007. The action potential in mammalian central neurons. Nature Reviews Neuroscience 8, 451–465.

[5] Becker, W., Kleinsmith, L., Bertoni, G., et al., 2009. Signal transduction mechanisms: I. electrical and synaptic signaling in neurons, in: The world of the cell. Benjamin Cummings, p.

[6] Biel, M., Wahl-Schott, C., Michalakis, S., Zong, X., 2009. Hyperpolarization-activated cation channels: from genes to function. Physiological reviews 89, 847–885.

[7] Breiman, L., 2001. Random forests. Machine learning 45, 5–32.

[8] Buettner, F., Natarajan, K.N., Casale, F.P., Proserpio, V., Scialdone, A., Theis, F.J., Teichmann, S.A., Marioni, J.C., Stegle, O., 2015. Computational analysis of cell-to-cell heterogeneity in single-cell rna-sequencing data reveals hidden subpopulations of cells. Nature biotechnology 33, 155–160.

[9] Cadwell, C.R., Palasantza, A., Jiang, X., Berens, P., Deng, Q., Yilmaz, M., Reimer, J., Shen, S., Bethge, M., Tolias, K.F., et al., 2016. Electrophysiological, transcriptomic and morphologic profiling of single neurons using patch-seq. Nature biotechnology 34, 199–203.

[10] Cadwell, C.R., Scala, F., Li, S., Livrizzi, G., Shen, S., Sandberg, R., Jiang, X., Tolias, A.S., 2017. Multimodal profiling of single-cell morphology, electrophysiology, and gene expression using patch-seq. Nature protocols 12, 2531–2553.

[11] Campanac, E., Gasselin, C., Baude, A., Rama, S., Ankri, N., Debanne, D., 2013. Enhanced intrinsic excitability in basket cells maintains excitatory-inhibitory balance in hippocampal circuits. Neuron 77, 712–722.

[12] Cardona-Alberich, A., Tourbez, M., Pearce, S.F., Sibley, C.R., 2021. Elucidating the cellular dynamics of the brain with single-cell rna sequencing. RNA biology 18, 1063–1084.

[13] Casale, A.E., Foust, A.J., Bal, T., McCormick, D.A., 2015. Cortical interneuron subtypes vary in their axonal action potential properties. Journal of Neuroscience 35, 15555–15567.

[14] Chamling, X., Kallman, A., Fang, W., Berlinicke, C.A., Mertz, J.L., Devkota, P., Pantoja, I.E.M., Smith, M.D., Ji, Z., Chang, C., et al., 2021. Single-cell transcriptomic reveals molecular diversity and developmental heterogeneity of human stem cell-derived oligodendrocyte lineage cells. Nature communications 12, 1–20.

[15] Consortium, T.U., 2022. UniProt: the Universal Protein Knowledge-base in 2023. Nucleic Acids Research 51, D523–D531.

[16] Darmanis, S., Sloan, S.A., Zhang, Y., Enge, M., Caneda, C., Shuer, L.M., Hayden Gephart, M.G., Barres, B.A., Quake, S.R., 2015. A survey of human brain transcriptome diversity at the single cell level. Proceedings of the National Academy of Sciences 112, 7285–7290.

[17] Dasgupta, D., Sikdar, S.K., 2019. Heterogeneous network dynamics in an excitatory-inhibitory network model by distinct intrinsic mechanisms in the fast spiking interneurons. Brain Research 1714, 27–44.

[18] Debanne, D., Inglebert, Y., Russier, M., 2019. Plasticity of intrinsic neuronal excitability. Current opinion in neurobiology 54, 73–82.

[19] Distasi, C., Dionisi, M., Ruffinatti, F.A., Gilardino, A., Bardini, R., Antoniotti, S., Catalano, F., Bassino, E., Munaron, L., Martra, G., et al., 2019. The interaction of sio2 nanoparticles with the neuronal cell membrane: activation of ionic channels and calcium influx. Nanomedicine 14, 575–594.

[20] Eze, U.C., Bhaduri, A., Haeussler, M., Nowakowski, T.J., Kriegstein, A.R., 2021. Single-cell atlas of early human brain development highlights heterogeneity of human neuroepithelial cells and early radial glia. Nature neuroscience 24, 584–594.

[21] Ge, S.X., Jung, D., Yao, R., 2020. Shinygo: a graphical gene-set enrichment tool for animals and plants. Bioinformatics 36, 2628–2629.

[22] Goldberg, E.M., Clark, B.D., Zagha, E., Nahmani, M., Erisir, A.,Rudy, B., 2008. K+ channels at the axon initial segment dampen near-threshold excitability of neocortical fast-spiking gabaergic interneurons. Neuron 58, 387–400.

[23] Golomb, D., Donner, K., Shacham, L., Shlosberg, D., Amitai, Y., Hansel, D., 2007. Mechanisms of firing patterns in fast-spiking cortical interneurons. PLoS Computational Biology 3, e156.

[24] Gouwens, N.W., Sorensen, S.A., Baftizadeh, F., Budzillo, A., Lee, B.R., Jarsky, T., Alfiler, L., Baker, K., Barkan, E., Berry, K., et al., 2020. Integrated morphoelectric and transcriptomic classification of cortical gabaergic cells. Cell 183, 935–953.

[25] Gouwens, N.W., Sorensen, S.A., Berg, J., Lee, C., Jarsky, T., Ting, J., Sunkin, S.M., Feng, D., Anastassiou, C.A., Barkan, E., et al., 2019. Classification of electrophysiological and morphological neuron types in the mouse visual cortex. Nature neuroscience 22, 1182–1195.

[26] Ha, G.E., Cheong, E., 2017. Spike frequency adaptation in neurons of the central nervous system. Experimental neurobiology 26, 179.

[27] Hao, Y., Hao, S., Andersen-Nissen, E., Mauck III, W.M., Zheng, S., Butler, A., Lee, M.J., Wilk, A.J., Darby, C., Zager, M., et al., 2021. Integrated analysis of multimodal single-cell data. Cell 184, 3573–3587.

[28] Herrera, A., Cheng, A., Mimitou, E.P., Seffens, A., George, D., Bar-Natan, M., Heguy, A., Ruggles, K.V., Scher, J.U., Hymes, K., et al., 2021. Multimodal single-cell analysis of cutaneous t-cell lymphoma reveals distinct subclonal tissue-dependent signatures. Blood, The Journal of the American Society of Hematology 138, 1456–1464.

[29] Hill, C.L., Stephens, G.J., 2021. An introduction to patch clamp recording. Patch Clamp Electrophysiology: Methods and Protocols, 1–19.

[30] Hodge, R.D., Bakken, T.E., Miller, J.A., Smith, K.A., Barkan, E.R., Graybuck, L.T., Close, J.L., Long, B., Johansen, N., Penn, O., et al., 2019. Conserved cell types with divergent features in human versus mouse cortex. Nature 573, 61–68.

[31] Hodgkin, A.L., Huxley, A.F., 1952. A quantitative description of membrane current and its application to conduction and excitation in nerve. The Journal of physiology 117, 500.

[32] Hwang, B., Lee, J.H., Bang, D., 2018. Single-cell rna sequencing technologies and bioinformatics pipelines. Experimental & molecular medicine 50, 1–14.

[33] Kim, H., Lee, J., Kang, K., Yoon, S., 2022. Markercount: A stable, count-based cell type identifier for single-cell rna-seq experiments. Computational and Structural Biotechnology Journal 20, 3120–3132.

[34] Kolb, B., Whishaw, I.Q., 1998. Brain plasticity and behavior. Annual review of psychology 49, 43–64.

[35] Kole, M.H., Letzkus, J.J., Stuart, G.J., 2007. Axon initial segment kv1 channels control axonal action potential waveform and synaptic efficacy. Neuron 55, 633–647.

[36] La Manno, G., Gyllborg, D., Codeluppi, S., Nishimura, K., Salto, C., Zeisel, A., Borm, L.E., Stott, S.R., Toledo, E.M., Villaescusa, J.C., et al., 2016. Molecular diversity of midbrain development in mouse, human, and stem cells. Cell 167, 566–580.

[37] Lake, B.B., Ai, R., Kaeser, G.E., Salathia, N.S., Yung, Y.C., Liu, R., Wildberg, A., Gao, D., Fung, H.L., Chen, S., et al., 2016. Neuronal subtypes and diversity revealed by single-nucleus rna sequencing of the human brain. Science 352, 1586–1590.

[38] Landau, S., Leese, M., Stahl, D., Everitt, B., 2011. Cluster Analysis. Wiley Series in Probability and Statistics, Wiley. URL: https://books.google.it/books?id=w3bE1kqd-48C.

[39] Lee, B., Dalley, R., Miller, J.A., Chartrand, T., Close, J., Mann, R., Mukora, A., Ng, L., Alfiler, L., Baker, K., et al., 2022. Signature morpho-electric properties of diverse gabaergic interneurons in the human neocortex. bioRxiv, 2022–11.

[40] Li, T., Tian, C., Scalmani, P., Frassoni, C., Mantegazza, M., Wang, Y., Yang, M., Wu, S., Shu, Y., 2014. Action potential initiation in neocortical inhibitory interneurons. PLoS biology 12, e1001944.

[41] Li, W., Luo, X., Ulbricht, Y., Wagner, M., Piorkowski, C., El-Armouche, A., Guan, K., 2019. Establishment of an automated patch-clamp platform for electrophysiological and pharmacological evaluation of hipsc-cms. Stem cell research 41, 101662.

[42] Lipovsek, M., Bardy, C., Cadwell, C.R., Hadley, K., Kobak, D., Tripathy, S.J., 2021. Patch-seq: Past, present, and future. Journal of Neuroscience 41, 937–946.

[43] Liu, H., Zhou, J., Tian, W., Luo, C., Bartlett, A., Aldridge, A., Lucero, J., Osteen, J.K., Nery, J.R., Chen, H., et al., 2021. Dna methylation atlas of the mouse brain at single-cell resolution. Nature 598, 120–128.

[44] Luecken, M.D., Theis, F.J., 2019. Current best practices in single-cell rna-seq analysis: a tutorial. Molecular systems biology 15, e8746.

[45] Marom, S., Marder, E., 2023. A biophysical perspective on the resilience of neuronal excitability across timescales. Nature Reviews Neuroscience, 1–13.

[46] Martini, L., Bardini, R., Savino, A., Di Carlo, S., 2022a. Gagam: a genomic annotation-based enrichment of scatac-seq data for gene activity matrix. bioRxiv.

[47] Martini, L., Bardini, R., Savino, A., Di Carlo, S., 2022b. Gagam v1.2: An improvement on peak labeling and genomic annotated gene activity matrix construction. Genes 14, 115.

[48] Martini, L., Bardini, R., Savino, A., Di Carlo, S., 2022c. High-resolution sample size enrichment of single-cell multi-modal low-throughput patch-seq datasets, in: 2022 IEEE International Conference on Bioinformatics and Biomedicine (BIBM), IEEE. pp. 2334–2341.

[49] Marx, V., 2022. Patch-seq takes neuroscience to a multimodal place. Nature Methods 19, 1340–1344.

[50] McFarlan, A.R., Chou, C.Y., Watanabe, A., Cherepacha, N., Haddad, M., Owens, H., Sjöström, P.J., 2022. The plasticitome of cortical interneurons. Nature Reviews Neuroscience, 1–18.

[51] Menon, S., Lui, V.C.H., Tam, P.K.H., 2021. Bioinformatics tools and methods to analyze single cell rna sequencing data. International Journal of Innovative Science and Research Technology,(IJISRT) 6, 282–288.

[52] Mi, H., Muruganujan, A., other, 2018. PANTHER version 14: more genomes, a new PANTHER GO-slim and improvements in enrichment analysis tools. Nucleic Acids Research 47, D419–D426. doi:10.1093/nar/gky1038.

[53] Morris, S.A., 2019. The evolving concept of cell identity in the single cell era. Development 146, dev169748.

[54] Mu, Q., Chen, Y., Wang, J., 2019. Deciphering brain complexity using single-cell sequencing. Genomics, Proteomics & Bioinformatics 17, 344–366.

[55] Mulas, C., Chaigne, A., Smith, A., Chalut, K.J., 2021. Cell state transitions: definitions and challenges. Development 148, dev199950.

[56] Neher, E., Sakmann, B., 1976. Single-channel currents recorded from membrane of denervated frog muscle fibres. Nature 260, 799–802.

[57] Neher, E., Sakmann, B., 1992. The patch clamp technique. Scientific American 266, 44–51.

[58] Obergrussberger, A., Friis, S., Brüggemann, A., Fertig, N., 2021. Automated patch clamp in drug discovery: major breakthroughs and innovation in the last decade. Expert opinion on drug discovery 16, 1–5.

[59] Papalexi, E., Satija, R., 2018. Single-cell rna sequencing to explore immune cell heterogeneity. Nature Reviews Immunology 18, 35–45.

[60] Platkiewicz, J., Brette, R., 2010. A threshold equation for action potential initiation. PLoS computational biology 6, e1000850.

[61] Poirion, O.B., Zhu, X., Ching, T., Garmire, L., 2016. Single-cell transcriptomics bioinformatics and computational challenges. Frontiers in genetics 7, 163.

[62] Potts, Y., Bekkers, J.M., 2022. Dopamine increases the intrinsic excitability of parvalbumin-expressing fast-spiking cells in the piriform cortex. Frontiers in Cellular Neuroscience 16, 919092.

[63] Rodríguez-Collado, A., Rueda, C., 2021. Electrophysiological and transcriptomic features reveal a circular taxonomy of cortical neurons. Frontiers in Human Neuroscience 15, 684950.

[64] Rogers, J., Gunn, S., 2005. Identifying feature relevance using a random forest, in: International Statistical and Optimization Perspectives Workshop” Subspace, Latent Structure and Feature Selection”, Springer. pp. 173–184.

[65] Rousseeuw, P.J., 1987. Silhouettes: a graphical aid to the interpretation and validation of cluster analysis. Journal of computational and applied mathematics 20, 53–65.

[66] Schober, P., Boer, C., Schwarte, L.A., 2018. Correlation coefficients: appropriate use and interpretation. Anesthesia & analgesia 126, 1763–1768.

[67] Seal, R.L., et al., 2022. Genenames.org: the HGNC resources in 2023. Nucleic Acids Research 51, D1003–D1009. doi:10.1093/nar/gkac888.

[68] Shore, A.N., Colombo, S., Tobin, W.F., Petri, S., Cullen, E.R., Dominguez, S., Bostick, C.D., Beaumont, M.A., Williams, D., Khodagholy, D., et al., 2020. Reduced gabaergic neuron excitability, altered synaptic connectivity, and seizures in a kcnt1 gain-of-function mouse model of childhood epilepsy. Cell reports 33.

[69] Spindler, A., Noble, S., Noble, D., 1999. Comparison of step and ramp voltage clamp on background currents in guinea-pig ventricular cells. Experimental physiology 84, 865–879.

[70] Stelzer, G., Rosen, N., Plaschkes, I., et al., 2016. The genecards suite: From gene data mining to disease genome sequence analyses. Current Protocols in Bioinformatics 54, 1.30.1–1.30.33. URL: https://currentprotocols.onlinelibrary.wiley.com/doi/abs/10.1002/cpbi.5, doi:10.1002/cpbi.5.

[71] Stephani, T., Hodapp, A., Jamshidi Idaji, M., Villringer, A., Nikulin, V.V., 2021. Neural excitability and sensory input determine intensity perception with opposing directions in initial cortical responses. Elife 10, e67838.

[72] Sugino, K., Clark, E., Schulmann, A., Shima, Y., Wang, L., Hunt, D.L., Hooks, B.M., Tränkner, D., Chandrashekar, J., Picard, S., et al., 2019. Mapping the transcriptional diversity of genetically and anatomically defined cell populations in the mouse brain. Elife 8, e38619.

[73] Szabó, A., Schlett, K., Szücs, A., 2021. Conventional measures of intrinsic excitability are poor estimators of neuronal activity under realistic synaptic inputs. PLoS Computational Biology 17, e1009378.

[74] Tepe, B., Hill, M.C., Pekarek, B.T., Hunt, P.J., Martin, T.J., Martin, J.F., Arenkiel, B.R., 2018. Single-cell rna-seq of mouse olfactory bulb reveals cellular heterogeneity and activity-dependent molecular census of adult-born neurons. Cell reports 25, 2689–2703.

[75] Tomar, R., 2019. Methods of firing rate estimation. BioSystems 183, 103980.

[76] Trojanowski, N.F., Bottorff, J., Turrigiano, G.G., 2021. Activity labeling in vivo using campari2 reveals intrinsic and synaptic differences between neurons with high and low firing rate set points. Neuron 109, 663–676.

[77] Turrigiano, G.G., Nelson, S.B., 2000. Hebb and homeostasis in neuronal plasticity. Current opinion in neurobiology 10, 358–364.

[78] Van Den Hurk, M., Bardy, C., 2019. Single-cell multimodal transcriptomics to study neuronal diversity in human stem cell-derived brain tissue and organoid models. Journal of Neuroscience Methods 325, 108350.

[79] Wang, M., Song, W.m., Ming, C., Wang, Q., Zhou, X., Xu, P., Krek, A., Yoon, Y., Ho, L., Orr, M.E., et al., 2022. Guidelines for bioinformatics of single-cell sequencing data analysis in alzheimer’s disease: review, recommendation, implementation and application. Molecular Neurodegeneration 17, 1–52.

[80] Welch, J.D., Kozareva, V., Ferreira, A., Vanderburg, C., Martin, C., Macosko, E.Z., 2019. Single-cell multi-omic integration compares and contrasts features of brain cell identity. Cell 177, 1873–1887.

[81] Wu, Y., Zhang, K., 2020. Tools for the analysis of high-dimensional single-cell rna sequencing data. Nature Reviews Nephrology 16, 408–421.

[82] Yang, J., Ye, M., Tian, C., Yang, M., Wang, Y., Shu, Y., 2013. Dopaminergic modulation of axonal potassium channels and action potential waveform in pyramidal neurons of prefrontal cortex. The Journal of physiology 591, 3233–3251.

[83] Yelhekar, T.D., Druzin, M., Karlsson, U., Blomqvist, E., Johansson, S., 2016. How to properly measure a current-voltage relation?—interpolation vs. ramp methods applied to studies of gabaa receptors. Frontiers in cellular neuroscience 10, 10.

[84] Zeng, H., 2022. What is a cell type and how to define it? Cell 185, 2739–2755.

